# Inference of continuous gene flow between species under misspecified models

**DOI:** 10.1101/2024.05.13.593926

**Authors:** Yuttapong Thawornwattana, Tomáš Flouri, James Mallet, Ziheng Yang

## Abstract

Gene flow between species is increasingly recognized as an important evolutionary process with potential adaptive consequences. Recent methodological advances make it possible to infer different modes of gene flow from genome-scale data, including pulse introgression at a specific time and continuous gene flow over an extended time period. However, it remains challenging to infer the history of species divergence and between-species gene flow from genomic sequence data. As a result, models used in real data analysis may often be misspecified, potentially leading to incorrect biological interpretations. Here, we characterize biases in parameter estimation under continuous migration models using a combination of asymptotic analysis and posterior inference from simulated datasets. When sequence data are generated under a pulse introgression model, isolation-with-initial-migration models assuming no recent gene flow are able to better recover gene flow with less bias than models that assume recent gene flow. When gene flow is assigned to an incorrect branch in the phylogeny, there may be large biases associated with the migration rate and species divergence times. When the direction of gene flow is incorrectly assumed, we may still detect gene flow if it is recent and between non-sister species but not when it is ancestral and between sister species. Overall, the impact of model misspecification is local in the species phylogeny. The pulse introgression model appears to be more robust to model misspecification and is preferable in real data analysis over the continuous migration model unless there is substantive evidence for continuous gene flow.

## Introduction

Gene flow between populations or species is an important process that shapes the genetic diversity we observe today. Genomic data can potentially revolutionize our understanding of the role of gene flow in adaptation and speciation. However, many methods to study gene flow from genomic data have largely been approximate (Hibbins and Hahn, 2022). They tend to focus on species triplets (or quartets if an an outgroup is used) and rely on summaries of sequence data, such as genome-wide site pattern counts (Green *et al*., 2010; Kubatko and Chifman, 2019), estimated gene tree topologies (Solis-Lemus and Ane, 2016; Jackson *et al*., 2017), or joint site frequency spectrum (Gutenkunst *et al*., 2009; Excofffier *et al*., 2021). These methods suffer from a loss of information and often have limited power to detect gene flow. Most methods cannot quantify the amount of gene flow or estimate key parameters such as effective population sizes and species divergence times (Jiao *et al*., 2021; Huang *et al*., 2022).

In this work, we focus on a class of methods that make use of the likelihood function calculated from genomic sequence data under the multispecies coalescent (MSC) model (Rannala and Yang, 2003), referred to as full-likelihood methods. Two idealized modes of gene flow have been modelled in the MSC framework. First, in the MSC-with-introgression (MSC-I) model (Flouri *et al*., 2020), also known as the multispecies network coalescent (MSNC; Yu *et al*., 2012; Wen *et al*., 2016; Wen and Nakhleh, 2018; Zhang *et al*., 2018) or network multispecies coalescent (NMSC) model (Ané *et al*., 2024), gene flow is assumed to occur in pulses at specific time points. The amount of gene flow is quantified by the introgression probability (φ_*A*→*B*_) from a source population *A* into a recipient *B*, which represents the proportion of migrants in *B* from *A* at the time of introgression (fig. 1**a**).

**Figure 1:**
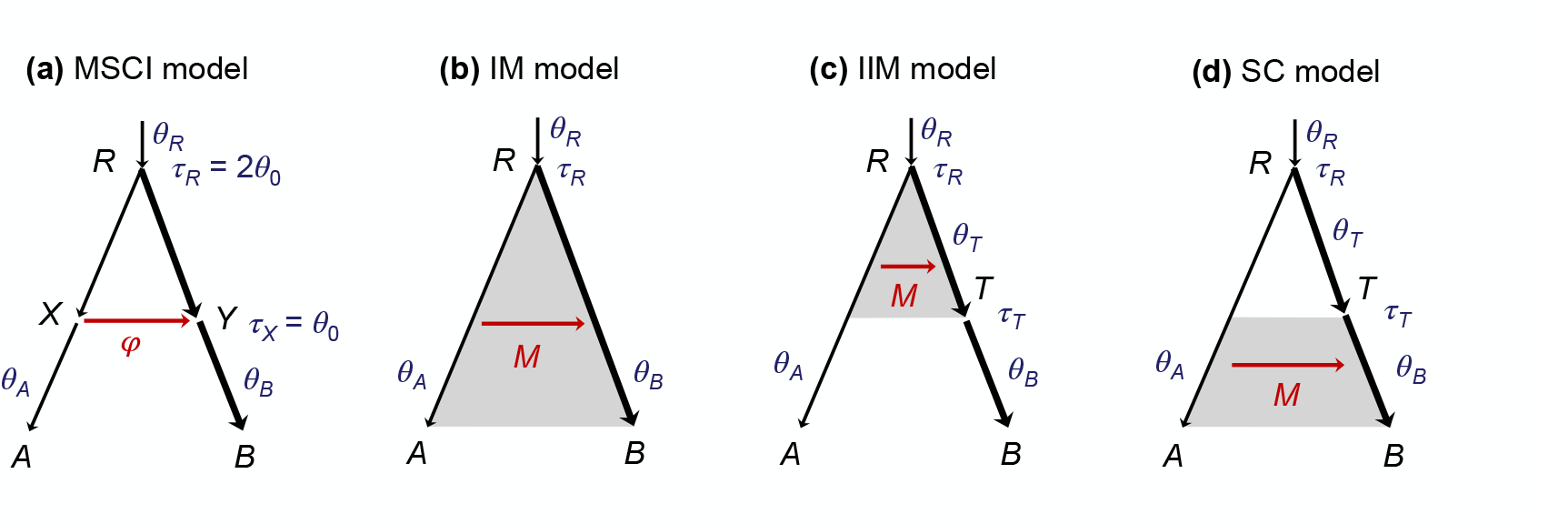
(**a**) MSC-I model for two species, *A* and *B*, used to generate data. Introgression occurs from *A* to *B* at time τ_*X*_ (forward in time) with probability φ. (**b**–**d**) Three MSC-M models used to analyze the data: IM (isolation with migration), IIM (isolation with initial migration), and SC (secondary contact). Each branch has an associated population size parameter *θ* = 4*Nµ*, where *N* is the effective population size and *µ* is the mutation rate per site per generation. Time is measured in the expected number of mutations per site, with τ = *T µ* where *T* is the divergence time in generations. In the MSC-I model (**a**), branch *X* has the same population size as *A*, and branch *Y* has the same population size as *B*. The parameter vector of the MSC-I model is Θ_i_ = {φ, τ_*X*_, τ_*R*_, *θ*_*X*_, *θ*_*R*_}. Grey shading indicates a period of continuous gene flow from *A* to *B* at rate *M*_*AB*_ = *N*_*B*_*m*_*AB*_ ≡ *M* migrants per generation where *m*_*AB*_ is the proportion of migrants in *B* from *A* per generation. In our numerical calculation and simulation studies, we use *θ*_*A*_ = *θ*_*R*_ = *θ*_0_ (thin branches), *θ*_*B*_ = 5*θ*_0_ (thick branches), τ_*X*_ = *θ*_0_, τ_*R*_ = 2*θ*_0_ and φ = 0.2, with *θ*_0_ = 0.002. The parameter vector of the IM model is Θ_IM_ = (τ_*R*_, *θ*_*A*_, *θ*_*B*_, *θ*_*R*_, *M*), while those for IIM and SC are Θ_IIM_ = Θ_SC_ = (τ_*R*_, τ_*T*_, *θ*_*A*_, *θ*_*B*_, *θ*_*T*_, *θ*_*R*_, *M*). The IIM and SC models are implemented in BPP as instances of the MSC-M model by including an unsampled ghost species that is sister to *B* and diverged with *B* at time τ_*T*_ . This creates two *θ* parameters for branches *RT* and *TB* as currently BPP does not implement the constraint *θ*_*T*_ = *θ*_*B*_.

Second, the MSC-with-migration (MSC-M) model (Flouri *et al*., 2023) assumes that gene flow occurs at a constant rate over an extended period of time. Gene flow from populations *A* to *B* is measured by the population migration rate *M*_*A*→*B*_ = *N*_*B*_*m*_*A*→*B*_, which is the expected number of individuals in *B* that are migrants from *A* per generation, *N*_*B*_ is the effective population size of *B*, and *m*_*A*→*B*_ is the proportion of individuals in *B* that have migrated from *A*. The MSC-M model is also known as the isolation-with-migration (IM) model (Nielsen and Wakeley, 2001; Hey and Nielsen, 2004; Hey, 2010; Zhu and Yang, 2012; Dalquen *et al*., 2017; Hey *et al*., 2018; Jones, 2019), although this often refers to the simplest instance of two species with migration occurring since their divergence up to the present (fig. 1**b**). Variants of MSC-M models include the isolation-with-initial-migration (IIM) model (fig. 1**c**; Costa and Wilkinson-Herbots, 2017) and the secondary contact (SC) model (fig. 1**d**; Costa and Wilkinson-Herbots, 2021).

Full likelihood implementations of these two modes of gene flow allow us to estimate key population parameters from genomic sequence data including the migration rates. The two modes, of course, represent extremes between which gene flow takes place in nature. Here, we use the term ‘introgression’ to refer to pulse gene flow in the MSC-I model while we use ‘migration’ strictly to refer to continuous gene flow in the MSC-M model. Both MSC models make the assumptions of no recombination within each locus and free recombination between loci; see Zhu *et al*. (2022) for an assessment of the impacts of intra-locus recombination on inference under the MSC.

Likelihood-based analysis of genomic sequence data requires one to specify a model of gene flow that best captures the system under study. There are many ways in which the assumed model may be misspecified. First, the mode of gene flow might be incorrectly assumed. For instance, gene flow might have occurred as a single pulse introgression event but the model assumes continuous gene flow. Second, lineages involved in gene flow may be incorrectly specified. This is particularly likely in large phylogenies where gene flow is detected in many species triplets and is assigned to ancestral branches using some heuristic criterion (Malinsky *et al*., 2018; Suvorov *et al*., 2022; Thawornwattana *et al*., 2023b; Ji *et al*., 2023). Currently, there is no simple method or principled way of assigning introgression to branches in a phylogeny. In theory, one could search over the space of MSC-I or MSC-M models in the Bayesian framework but this task remains computationally intractable except for very small datasets (Wen and Nakhleh, 2018; Zhang *et al*., 2018). Third, the source and recipient populations of gene flow may be incorrectly specified: most summary methods cannot identify the direction of gene flow. When the model of gene flow is misspecified, the resulting inferences can be biased, potentially leading to erroneous biological interpretations. It is thus important to understand the nature of the biases induced by model misspecification, and the conditions under which certain important parameters may still be reliably estimated.

Here, we investigate the impact of these three kinds of model misspecification on Bayesian parameter estimation under the MSC-M model. We assume continuous migration in the model but gene flow may actually occur as pulse introgression (as assumed in the MSC-I model), or in the opposite direction, or involve wrong branches on the species phylogeny that are parental or daughter to the lineage genuinely involved in gene flow. We characterize the bias and variance of parameter estimates using a combination of asymptotic analysis and posterior inference of simulated mutilocus datasets under different misspecification scenarios. We are particularly interested in the impact on the estimates of species divergence times and population sizes, as well as how estimates of the migration rate (*M*) corresponds to the introgression probability (φ) in the data-generating model. We use the Bayesian program BPP (Flouri *et al*., 2018, 2023) to analyze the data but our results should apply to other full-likelihood methods (Wen and Nakhleh, 2018; Zhang *et al*., 2018). This work complements our previous studies on model misspecification when the MSC-M is assumed to be true and data are analyzed under the MSC-I model (Jiao *et al*., 2020; Huang *et al*., 2022) and when the direction of introgression was misspecified under the MSC-I model (Thawornwattana *et al*., 2023a).

## Results

### Correspondence between the MSC-I and MSC-M models: the case of two species

If gene flow occurs from *A* to *B* at time τ _*X*_ with probability φ but we analyze the data using a model that assumes continuous migration over a certain time period to fit the sequence data, what will the estimate of the migration rate *M* be like? Will we detect gene flow despite misspecification of the mode of gene flow? Similarly, what are the effects of misspecified direction of gene flow? We approach these questions using a combination of asymptotic analysis as the number of loci approaches infinity and Bayesian inference of simulated sequence data, following Jiao *et al*. (2020) and Huang *et al*. (2022). The asymptotic analysis is only tractable in the simplest case of two species with one sequence per species. By contrast, simulations can be performed for any number of species, any number of sequences per species, and any finite number of loci. In this work, we generated sequence data under the MSC-I model with parameters Θ_i_ (e.g., fig. 1**a**). We are interested in quantifying the biases in the estimates of parameters Θ_m_ under the MSC-M model (e.g., fig. **1b-d**).

We first approach this problem using the asymptotic theory that as the number of loci *L* → ∞, the maximum likelihood estimate (MLE) 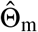 of Θ_m_ under the MSC-M model converges to 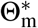 which minimizes the Kullback–Leibler (KL) divergence

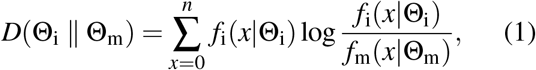

where *f*_i_(*x*|Θ_i_) is the probability of observing *x* differences (at *n* sites) under the true MSC-I model and represents the data, and where *f*_m_(*x*|Θ_m_) is the corresponding probability under the fitting MSC-M model (Jiao *et al*., 2020; Huang *et al*., 2022). Because of model mismatch, a perfect fit is impossible, with 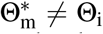, but 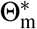 is the best-fitting parameter value under the MSC-M model.

We consider the simplest case of two species, *A* and *B* (fig. 1), with the data consisting of an infinite number of loci of length *n*. Each locus has one sequence from each species, *a* and *b*, summarized as *x* differences out of *n* sites. The probability for *x* is given by averaging over the unobserved coalescent time *t* between the two sequences,

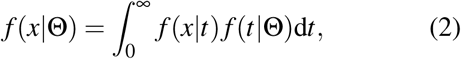

where the density of coalescent time, *f* (*t*|Θ), depends on the models of gene flow, and is given in SI text for the MSC-I model of figure 1**a** and the MSC-M models of figure **1b-d**, and where the probability for the number of nucleotide differences given the coalescent time, *f* (*x*|*t*), is given by the mutation model.

For analytical tractability, we assume the infinite-sites mutation model instead of a finite-sites model such as JC (Jukes and Cantor, 1969), so that *f* (*x*|*t*) is given by the Poisson probability as

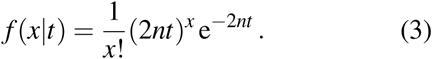

This leads to a closed-form expression of *f* (*x*|Θ) as in Huang *et al*. (2022).

Of particular interest is the correspondence between the introgression probability φ in the MSC-I model and the migration rate *M* in the MSC-M model. Under the MSC-M model, the probability that a lineage from species *B* traces back to *A* is

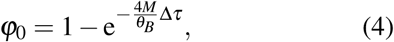

where Δ_τ_ is the time period of gene flow (Huang *et al*., 2022). Inverting this relationship gives

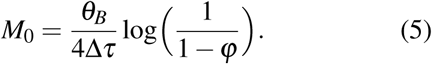

When φ is small, 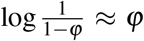, so that 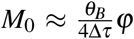 or 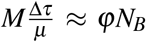: the total number of migrants over the time period of Δ τ */µ* generations under the MSC-M model is approximately equal to the number of migrants under the MSC-I model. Eq. 5 gives the expected migration rate when the true model is MSC-I with introgression probability φ under the assumption that all gene flow that has occurred is ‘recovered’ by the fitting MSC-M model.

#### Asymptotic analysis of MLEs

We now study the asymptotic behavior of parameter estimation under the migration (MSC-M) models of figure **1b-d** when the data are generated under the introgression (MSC-I) model (fig. 1**a**) in the case of two species. With one sequence per species per locus, the parameter vector for the true MSC-I model is Θ_i_ = { φ, *τ* _*X*_, *τ* _*R*_, *θ*_*X*_, *θ*_*R*_}, while those for the fitting MSC-M models are Θ_IM_ = {*M, τ* _*R*_, *θ*_*A*_, *θ*_*R*_} and Θ_IIM_ = Θ_SC_ = {*M, τ* _*T*_, *τ* _*R*_, *θ*_*A*_, *θ*_*R*_} (fig. 1**b-d**). Using eq. 1, we obtain the asymptotic MLEs 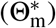 under the IM, IIM and SC models (fig. 2) for data of an infinite number of loci. We assume *θ*_*X*_ = *θ*_*R*_ in the MSC-I model and *θ*_*A*_ = *θ*_*R*_ in the fitting MSC-M model. The true and best-fitting distributions of the coalescent time *t*_*ab*_ = *t* are shown in figure S1. We varied the sequence length (*n*) and the introgression probability (φ).

**Figure 2:**
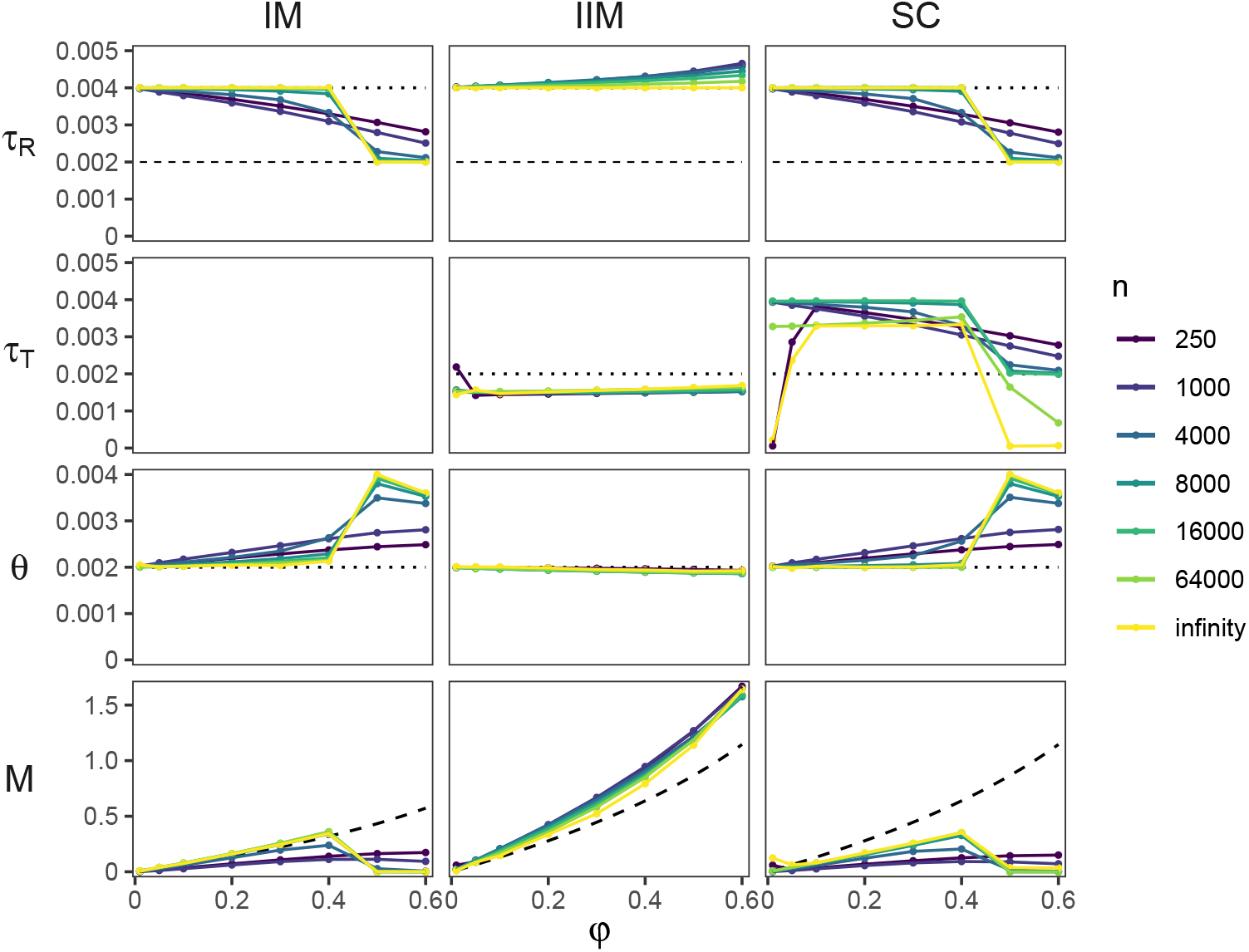
Best-fitting parameter values under the IM, IIM and SC models (fig. 1**b**–**d**) when the data consist of one sequence (of *n* sites) per species generated under the MSC-I model (fig. 1**a**). The two population sizes, *θ*_*A*_ and *θ*_*R*_, were constrained to be equal, denoted by *θ* . Horizontal dotted lines indicate true values. For τ_*R*_, the dashed line indicates the introgression time τ_*X*_ in the MSC-I model. For *M*, dashed curves indicate the expected value *M*_0_ based on eq. 5, assuming the true *θ*_*B*_ and the expected duration of migration to be Δτ = τ_*R*_ for the IM model and Δτ = τ_*R*_ − τ_*X*_ for the IIM and SC models. The true and best-fitting distributions of the coalescent time (*t*) are in figure S1.

Among the three MSC-M models of figure 1**b-d**, the IIM model provide the most sensible parameter values, with the best-fitting migration rate *M*^∗^ tracking to the introgression probability (φ) in the MSC-I model (fig. 2). The best-fitting 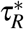 and 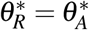 also match the true values. This means coalescence in the ancestral population (*R*) is correctly accounted for. The time at which migration stops 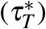 is slightly younger than the actual introgression time 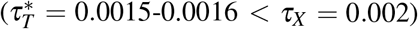 due to a mismatch in the mode of gene flow. Under the MSC-I model, gene flow results in a peak in the density of the coalescent time *t*_*ab*_ at τ _*X*_ (fig. S1, black curve). By contrast, coalescence due to migration under the IIM model peaks in the middle of the migration period (fig. S1, dark blue curve, middle column). Thus having 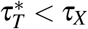 gives a better fit.

The IM and SC models show similar patterns of best-fitting parameter values, which are distinct from the IIM model. Under the IM model, the absence of recent gene flow in the MSC-I model (fig. 1**a**) leads to *M*^∗^ being much smaller than that from the IIM model (fig. 2, first column). At small values of 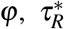 matches the true value as the number of sites approaches infinity, and the estimated migration rate matches the expected value (*M*_0_) from eq. 5 assuming the migration period (0, τ _*R*_) and the true population size *θ*_*B*_ (fig. 2, first column). At large values of φ (say φ *>* 0.5), 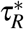 drops to τ _*X*_ (figs. 2 and S1). This is coupled with a drop in *M*^∗^ and an overestimation of *θ*∗. Thus as the introgression probability becomes large, it becomes increasingly more difficult for the IM model to explain the absence of recent gene flow in the MSC-I model because increasing the value of *M*^∗^ would worsen the fit (fig. S1). Instead, the IM model prefers a scenario with a recent divergence time 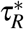 and a very low migration rate *M*^∗^ after the divergence. In this case, all variation variation due to introgression in the data-generating model is explained by coalescence in the common ancestor. Shorter loci provide weaker evidence of gene flow since similar sequences can be explained by random mutations and more recent divergence. As a result, we obtained smaller *M*^∗^, younger 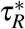, and larger *θ*∗. We explain the effect of sequence length in more detail in the next section. Finally, under the SC model, the time at which migration starts 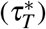 is often close to the divergence time 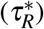, making it similar to the IM model.

#### Simulation studies

Our asymptotic analysis assumes two sequences sampled per locus (with one sequence per species). To study more complex settings, we used BPP to analyze simulated datasets. The data were simulated under the MSC-I model (fig. 1**a**) and analyzed under the three MSC-M models of figure 1**b-d**. The implementation in BPP allows an arbitrary number of species and sequences per species per locus, and the likelihood calculation averages over the gene genealogy underlying the sequence alignment at each locus. We assume the JC mutation model (Jukes and Cantor, 1969). In the base case, we set the introgression probability φ = 0.2 and each dataset consists of *L* = 4,000 loci, with *S* = 4 sequences sampled per species per locus, and each sequence has *n* = 1,000 sites. We varied the following four factors to examine their impact on posterior estimates of parameters, in particular the migration rate *M*: the number of sites per sequence (*n*), the number of sequences per species (*S*), the number of loci per species (*L*), and the introgression probability (φ). The first three factors are related to data size while the last is a parameter that measures the strength of gene flow. The results are summarized in figure 3.

**Figure 3:**
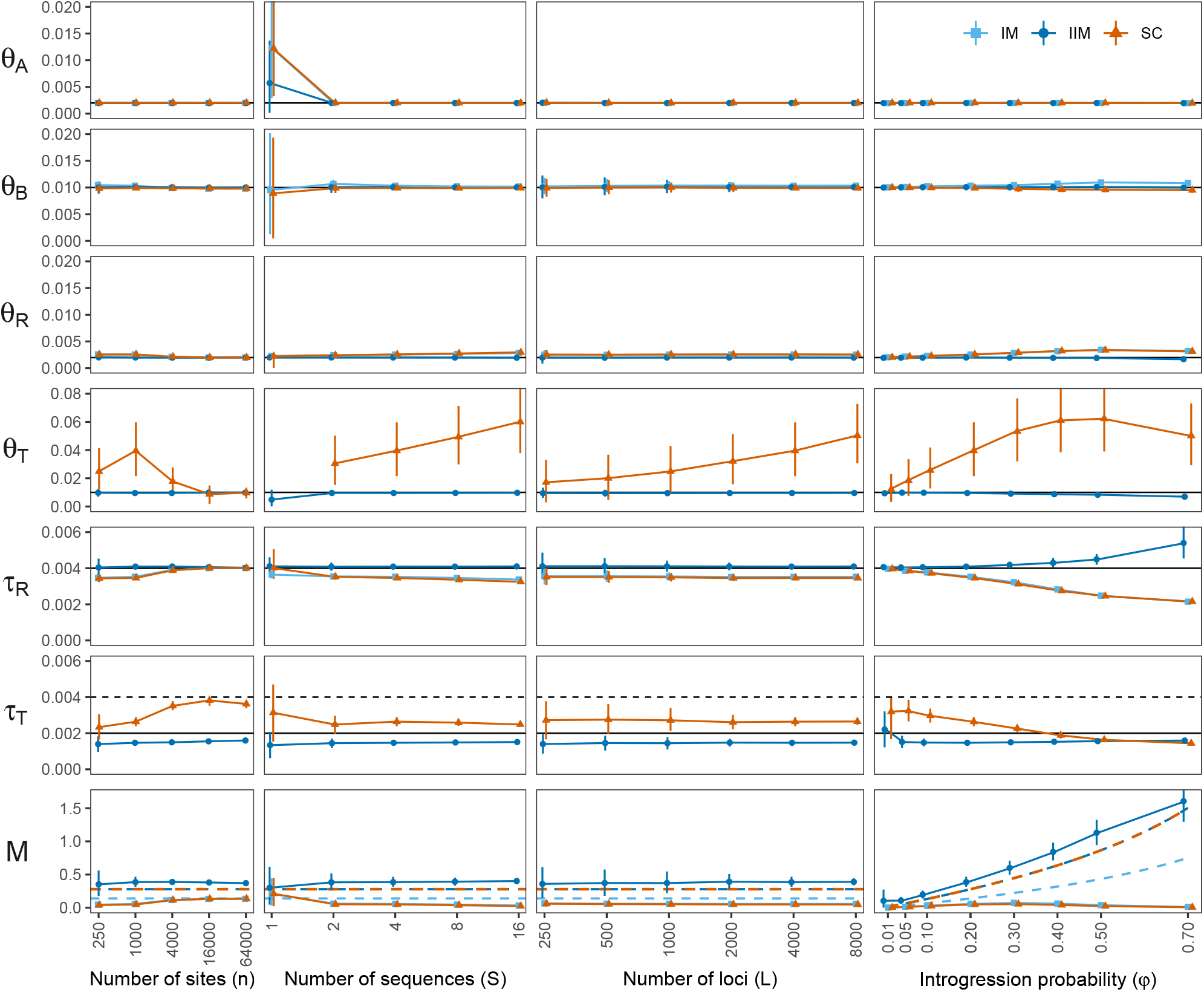
Parameter estimates from the three MSC-M models (IM, IIM and SC; fig. 1**b-d**) of data generated under the MSC-I model (fig. 1**a**), summarized as posterior mean and 95% HPD CIs averaged over 30 replicate datasets. In the base case, each dataset consists of *L* = 4,000 loci, with *S* = 4 sequences per species at each locus and *n* = 1,000 sites per sequence. Parameters in the MSC-I model are given in the legend to figure 1. We varied four factors one at a time, keeping other factors fixed at the base case: the number of sites per sequence (*n*), the number of sequences per species (*S*), the number of loci (*L*), and the introgression probability (φ). The parameters *θ*_*T*_ and τ_*T*_ are specific to the IIM and SC models. When *S* = 1, *θ*_*B*_ is unidentifiable in the IIM model (fig. 1**c**), and *θ*_*T*_ is unidentifiable in the SC model (fig. 1**d**). Horizontal solid lines indicate the true values used to generate the data. For τ_*T*_, the additional horizontal dotted line indicates τ_*R*_, its upper limit. For *M*, a dashed curve indicates the expected value *M*_0_ based on eq. 5, assuming the true *θ*_*B*_ and the expected duration of migration Δτ; see legend to figure 2. The *x*-axis for *n, S* and *L* is on the logarithmic scale.

First, we examined the effects of sequence length *n* (fig. 3, first column). This has little impact on the posterior mean and highest-probability-density (HPD) credibility intervals (CIs) for *θ*_*A*_, *θ*_*B*_, *θ*_*R*_, and τ_*R*_ under all three models, as well as *θ*_*T*_ and τ_*T*_ under the IIM model. Even with short sequences (*n* = 250), those parameters were precisely estimated. However, use of longer sequences improves the precision in the estimated migration rate *M* under the IIM model.

At *n* = 64,000 sites, the estimate 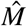 is 0.37, which is close to the limiting value *M*^∗^ = 0.33 from our asymptotic analysis of data of infinitely many loci of two sequences under the assumption of equal population sizes. When the IM or SC model was used to fit the data, we obtain much smaller estimates of *M*, again agreeing with our asymptotic results (fig. 2).

We also conducted the Bayesian test of gene flow, comparing the null hypothesis *H*_0_ : *M*_*A*→*B*_ = 0 against the alternative hypothesis *H*_1_ : *M*_*A*→*B*_ *>* 0 (fig. S3**a**).

The test has high power, rejecting the null and supporting gene flow with the Bayes factor *B*_10_ *>* 100 in nearly all datasets. Interestingly, for the IM and SC models, 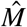 increases from 0.05 at *n* = 250 to 0.13 at *n* = 64,000. This trend is accompanied by an increase in 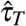 in the SC model, which converges to τ_*R*_ at large values of *n* (fig. 3; also see fig. S2). Thus the SC model becomes equivalent to the IM model in the limit as *n* → ∞. This pattern of increasing intensity 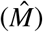 and duration 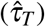 of gene flow in the SC model reflects the increasingly stronger evidence of gene flow in longer sequences. In the MSC-I model, introgression leads to a reduction in the coalescent time *t*_*ab*_. When sequences are short, this small *t*_*ab*_ can be accounted for in the fitting model as random mutations or recent divergence leading to nearly identical sequences without having gene flow. This explains small values of 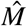 and an underestimation of τ_*R*_ when loci are short (fig. 3; *n* = 250 and 1,000). As sequences become longer, the chance that they are similar purely due to mutations becomes increasingly low. Thus increasing the migration rate can reduce *t*_*ab*_ and make the data seem more plausible. This also improves 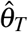 because migration helps explain large genetic variation in the recipient population *B* as a result of introgression from *A* in the MSC-I model, and the coalescent time *t*_*bb*_ of sequences from *B* shows an improved fit (fig. S2).

When 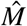 is low (e.g., at *n* = 250 or 1,000), large values of 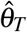 are needed to explain this variation. Finally, 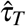 under the IIM model is estimated to be about 0.0015, as predicted by our asymptotic analysis of both finite and infinitely-long sequences (fig. 2), while the true introgression time τ_*X*_ is 0.002. One might expect 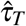 (the time at which the migration period ends) to converge to τ_*X*_ since this is the smallest time at which sequences from *A* and *B* can coalesce (i.e., the smallest *t*_*ab*_). We obtain 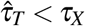. This may be partly attributable to the difference in how the MSC-I and the IM models account for the reduced *t*_*ab*_ due to gene flow. The probability density of *t*_*ab*_ peaks at the introgression time τ_*X*_ in the MSC-I model while it peaks in the middle of the migration period (τ_*T*_, τ_*R*_) in the IIM model (see fig. S1). Thus having a migration period that ends after τ_*X*_ better accommodates *t*_*ab*_ in the data. In summary, the IIM model is able to detect more gene flow and provides more precise and accurate estimates of population sizes and divergence times even with short sequences (*n* = 250) while the IM and SC models require at least 4,000 sites per locus to be able to detect good amounts of gene flow (fig. 3; 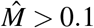).

Next, we consider the effects of the number of sequences per species (*S*). When there is only one sequence from each species in the data (*S* = 1), no coalescent events can occur in *B* and *T* . Thus *θ*_*B*_ is not identifiable under the IIM model, and *θ*_*T*_ is not identifiable under the SC model. In other cases (*S >* 1), all parameters in each of the MSC-M models (Θ_IM_, Θ_IIM_, Θ_SC_) are identifiable. When *S* = 1, estimates of *θ*_*A*_, *θ*_*B*_ and *M* have wide CIs in all three fitting models (note that there is no *θ*_*B*_ in the IIM model in this case), with *θ*_*A*_ being grossly overestimated (fig. 3, second column). There is a poor fir to the coalescent time *t*_*aa*_ for sequences from *A* (fig. S4). Nonetheless, the ancestral population size *θ*_*R*_ is well-estimated. When there were at least two sequences from each species (*S* ≥ 2), all parameters are estimated with narrow CIs, except for *θ*_*T*_ in the SC model for the same reason as in the case of varying *n*. As before, the IIM model is able to infer migration, with 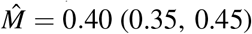 at *S* = 16. The IM and SC models detect much less gene flow, with 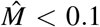, although gene flow detected is always significant (fig. S3). The values of 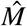 decreases slightly as *S* increases. This trend is correlated with a slight decrease in 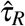, an increase in 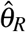, and a large increase in 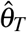 (fig. 3).

Third, the effect of the number of loci (*L*) was similar to that of *S* (figs. 3, third column and S5). Using a small number of loci (*L <* 1,000) results in estimates of *θ*_*B*_, *M* and *θ*_*T*_ having wide CIs. In theory, the CI width should reduce by a half as *L* (dataset size) quadruples. This holds approximately for most parameters except for *θ*_*T*_ in the SC model, which is poorly estimated.

Lastly, we consider the impact of the introgression probability (φ) in the MSC-I model. For all fitting models, the extant (*θ*_*A*_, *θ*_*B*_) and ancestral (*θ*_*R*_) population sizes are well-estimated, with the posterior mean close to the true value and with narrow CIs. Consistent with our asymptotic results, only the IIM model is able to estimate *M* that increases with φ (fig. 3, last column). As φ increases, the divergence time τ_*R*_ becomes increasingly overestimated while 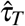 stays largely unchanged, resulting an increasingly long period of migration 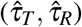. This means deep coalescent events between sequences from *A* and *B* in *R* are being interpreted as a result of migration after species divergence (fig. 4). By contrast, the IM and SC models only detect small amounts of gene flow (*M <* 0.1) regardless of the true value of φ (fig. 3). At extreme values of φ, gene flow detected by the IM or SC model is often not significant (fig. S3**d**). In the MSC-I model, larger values of φ lead to smaller *t*_*ab*_, with a peak at the introgression time (τ_*X*_ ). The IM and SC models accommodate small *t*_*ab*_ in the data as coalescence in the ancestral population, with 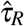 gradually decreasing from τ_*R*_ to τ_*X*_ as φ increases. This reduction in 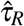 is associated with an increase in 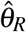. This pattern agrees with our asymptotic analysis (fig. 2, *n* = 1,000). It also predicts that 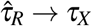 as *n* → ∞ and *L* → ∞. Lastly, one difference between the IM and SC models is in the estimate of *θ*_*B*_. As φ increases, there is a slight overestimation of *θ*_*B*_ under the IM model but an underestimation under the SC model (fig. 3).

**Figure 4:**
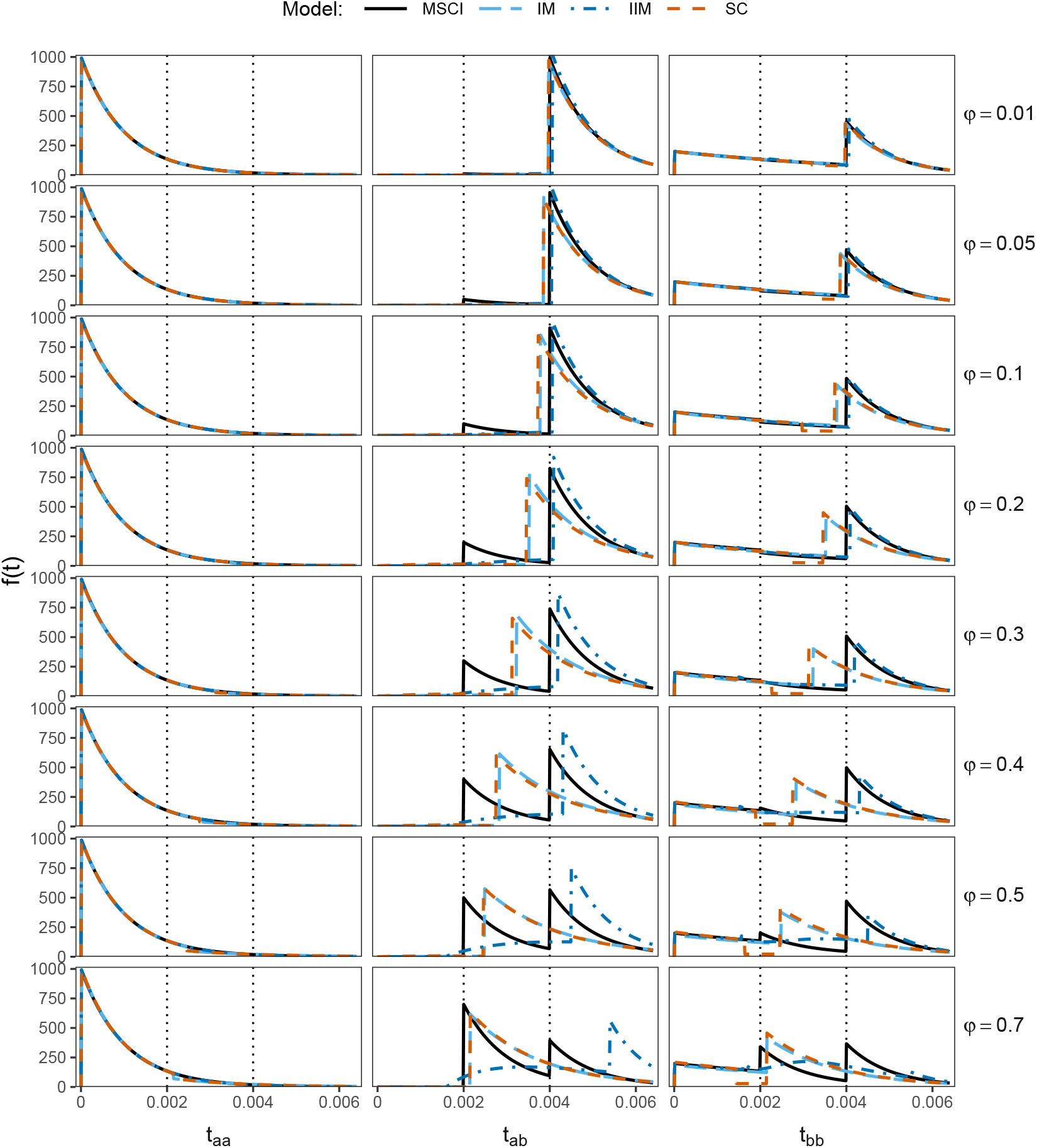
Distributions of coalescent times between two sequences (*t*_*aa*_, *t*_*ab*_ and *t*_*bb*_) under the true MSC-I model (fig. 1**a**) at different values of the introgression probability (φ; solid curves) and under the three fitting MSC-M models (IM, IIM, SC in fig. 1**b-d**; dashed curves) calculated using the approximate best-fitting parameter values obtained from BPP for data of *L* = 4,000 loci, each with *S* = 4 sequences per species and with *n* = 1,000 sites per sequence (fig. 3, last column). The two vertical dotted lines indicate the introgression time (τ_*X*_ = 0.002) and the divergence time (τ_*R*_ = 0.004).

Why do the IM and SC models detect much less gene flow than the IIM model across a wide range of values of φ? In these two models, migration stops at the present. Thus estimating some positive values of *M* means predicting coalescence events between *A* and *B* during (0,τ_*X*_ ) while none exist in the data-generating model. This scenario is thus a worse fit than detecting no gene flow at all.

When the SC model is used to fit the data, 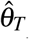 is usually much larger than the true value and with a wide CI, and 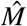 is close to zero. This is because under the MSC-I model, introgression from *A* into *B* leads to increased genetic variation in *B*. Here, with *θ*_*B*_ being well-estimated and *M* being close to zero, having a large value of 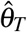 helps explain genetic variation in *B*. This also explains why 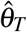 peaks at φ = 0.5, where genetic variation in *B* is the largest under the MSC-I model (fig. 3).

Overall, these simulation results agree with our asymptotic analysis of one sequence per species assuming an infinite number of loci (fig. 2; *n* = 1,000). Among the three MSC-M models, the IIM model provides the most sensible estimates and is able to estimate corresponding amounts of gene flow in the data generated under the MSC-I model. However, the divergence time (τ_*R*_) is overestimated when the introgression probability (φ) is high. The IM and SC models are qualitatively similar: they can recover much less gene flow than the IIM model and require long sequences of at least *n* = 4,000 sites per locus to detect a reasonable amount of gene flow. We find no major impacts on parameter estimates of *n, S*, and *L*, all of which reflect the dataset size.

### The case of four species

We extend our simulation study to more complex cases involving four species with the phylogeny ((*A*, (*B, C*)), *D*) and with gene flow between *A* and *B* or between *A* and the common ancestor of *B* and *C* (fig. 5). Species *D* serves as an outgroup and is not involved in gene flow. We simulated data under models A, B, C, or D and analyze them under models C and D (fig. 5). Gene flow may thus be misspecified in two ways. First, migration may be assigned to a wrong branch (C-D and D-C settings). Second, the mode of gene flow is incorrectly assumed (A-C and B-D settings). We also consider a combination of both kinds of misspecification (B-C and A-D settings). Note that model C is an instance of the SC model while model D is an instance of the IIM model, and that gene flow is between non-sister lineages in model C and between sister species under mode D. We simulated datasets consisting of *L* = 250, 1,000, or 4,000 loci, with *S* = 4 sequences per species per locus and with each sequence having *n* = 500 sites. The results are summarized in figure 5 and table S1.

**Figure 5:**
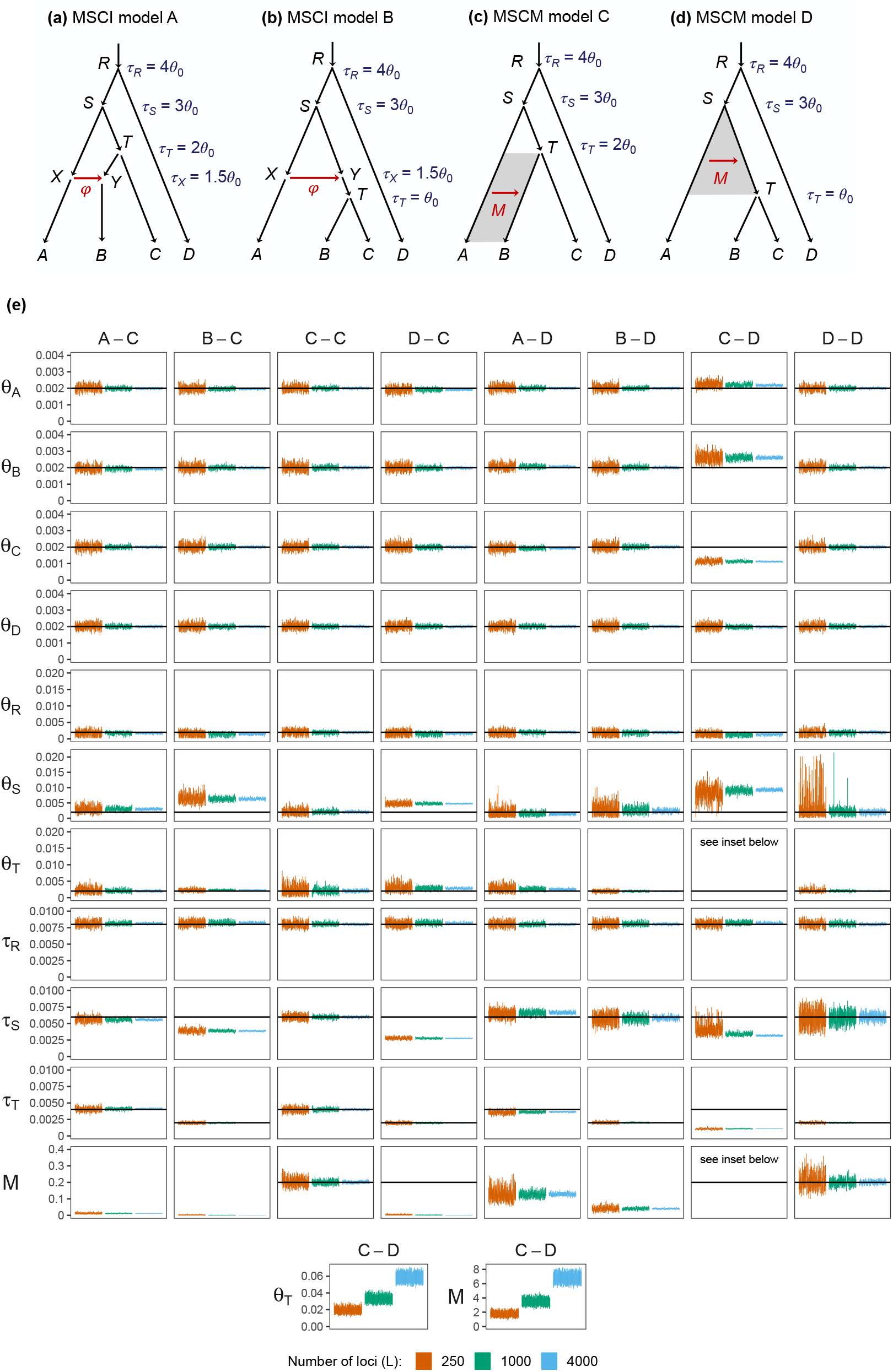
(**a-b**) Two introgression (MSC-I) models and (**c-d**) two migration (MSC-M) models used in simulation. All branches have population size *θ*_0_ = 0.002. In MSC-I model A, the species divergence and introgression times are τ_*R*_ = 4*θ*_0_, τ_*S*_ = 3*θ*_0_, τ_*T*_ = 2*θ*_0_, and τ_*X*_ = τ_*Y*_ = 1.5*θ*_0_. In MSC-I model B, τ_*R*_ = 4*θ*_0_, τ_*S*_ = 3*θ*_0_, τ_*T*_ = *θ*_0_, and τ_*X*_ = τ_*Y*_ = 1.5*θ*_0_. Introgression probability is φ = 0.2. In MSC-M model C, τ_*R*_ = 4*θ*_0_, τ_*S*_ = 3*θ*_0_, and τ_*T*_ = 2*θ*_0_, with migration occurring from species *A* to *B* during (0, τ_*T*_ ) at rate *M* = 0.2 migrants per generation. In MSC-M model D, τ_*R*_ = 4*θ*_0_, τ_*S*_ = 3*θ*_0_, and τ_*T*_ = *θ*_0_, with migration from *A* to *T* during (τ_*T*_, τ_*S*_) at rate *M* = 0.2. (**e**) The 95% HPD CIs of parameters (rows) from 100 replicate datasets of *L* = 250, 1,000, and 4,000 loci. Column labels refer to the simulation model followed by the analysis model; e.g., ‘A-C’ means the data were simulated under model A and analyzed under model C. Black solid line indicates the true value. Estimates of *θ*_*T*_ and *M* for the C-D setting are shown separately at the bottom due to their extreme values.

The four models (A-D) used here are the same as used in Huang *et al*. (2022), where complementary results from using the MSC-I model to analyze data generated under the MSC-M model can be found.

#### Inference under the correct model (fig. 5, C-C and D-D)

As a reference for comparison, we first examine the two cases in which the model used to analyze the data is correct: C-C and D-D (fig. 5**e**). In both settings, most parameters, including the migration rate *M* (see table S1), are well estimated, with a decreasing CI width as the number of loci (*L*) increases. Parameters that are more difficult to estimate are *θ*_*R*_ in the C-C setting and *θ*_*R*_, *θ*_*S*_ and τ_*S*_ in the D-D setting. Their estimates have considerably wider CIs even when *L* = 4,000 loci were used.

#### Migration assigned to a wrong branch (fig. 5, C-D and D-C)

In the C-D setting, the data were generated under model C, with migration occurring after a period of isolation, but were analyzed under model D, with migration occurring right after species divergence. We find that only the root divergence time (τ_*R*_) and the outgroup population size (*θ*_*D*_) are well estimated. Under the data-generating model C, sequences from *A* are expected to be closer to those from *B* than to those from *C* due to migration, with *t*_*ab*_ *< t*_*ac*_.

However under the fitting model D, those sequences should be equidistant, with *t*_*ab*_ = *t*_*ac*_. Thus there is a conflict between the data-generating and fitting models in explaining those coalescent times (fig. S6). Under model D, the divergence times τ_*T*_ and τ_*S*_ are thus severely underestimated to accommodate the small *t*_*ab*_ in the data, with 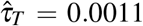 and 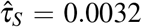 while the true values are τ_*T*_ = 0.002 and τ_*S*_ = 0.006. With τ_*S*_ underestimated, *θ*_*S*_ is overestimated. The estimates of *θ*_*B*_ and *θ*_*C*_ are also affected, with 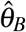 overestimated and 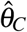 underestimated. Another conflict is that in the fitting model D, *t*_*ab*_ and *t*_*ac*_ have the same distribution while they are different under the data-generating model C (fig. S6, third row). This leads to a poor fit of *t*_*ac*_. Estimates of the migration rate and recipient population size (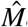 *and*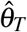 ) are unreasonably large. However, their ratio or the mutation-scaled migration rate *M/θ*_*T*_ = *m/µ* is much better estimated, with 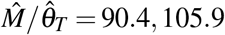, and 116.3 for *L* = 250, 1,000 and 4,000, respectively, compared with the true value in model C of *M/θ*_*T*_ = 100 (table S1). This suggests that the proportion of migrants (*m*) has a greater impact on the distribution of gene trees and coalescent times than the number of migrants (*M* = *Nm*).

In the D-C setting, migration occurs initially after species divergence (i.e., IIM) but the analysis model assumes secondary contact (SC). Population sizes of the modern species (*θ*_*A*_, *θ*_*B*_, *θ*_*C*_, *θ*_*D*_) are very well estimated, as are the population size and age of the root (*θ*_*R*_, τ_*R*_) and the divergence time 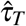 between *B* and *C*. However, τ_*S*_ and *θ*_*S*_ are grossly biased (fig. 5**e**). Since *B* is the recipient of migration from *A* in the fitting model D, the estimated migration rate is very low (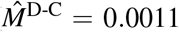 ; table S1) and is not significant (fig. S7). As a result, recent coalescent events between *A* and *T* in the data lead to an excess of the coalescent times *t*_*ab*_ and *t*_*ac*_, and a deficit in *t*_*bc*_ during (τ_*T*_, τ_*S*_) (fig. S6). In the fitting model D, this pattern is explained by having a very recent divergence time τ_*S*_ between *A* and *T*, which is estimated to be close to 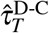, with 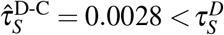 and 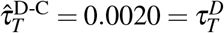 (table S1). Because of this underestimation of τ_*S*_, *θ*_*S*_ is overestimated. Overall, the fitted distribution of *t*_*bc*_ matches the true distribution under model C reasonably well while the fitted values of *t*_*ab*_ and *t*_*ac*_ reflect the increased duration 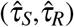 of population *S*, with the majority of coalescent events occurring closer to 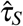 (fig. S6).

In summary, assigning a migration event incorrectly onto parental or daughter branches to the lineages genuinely involved in gene flow can have devastating effects on estimation of the rate of gene flow and of related divergence-time and population-size parameters. Nevertheless, the impact of misspecification is local, affecting only parameters related to the parts of the species phylogeny.

#### Wrong mode of gene flow: continuous migration versus pulse introgression (fig. 5, A-C and B-D)

Next, we consider cases where the mode of gene flow is misspecified but the population pair involved in gene flow is correctly specified, with data generated under MSC-I and analyzed under MSC-M. In the A-C setting, gene flow is between non-sister species while in the B-D setting, it is between sister species. This is an extension of our two species analysis (figs. 1–4) to a larger phylogeny.

In both A-C and B-D settings, all population size and divergence time parameters are correctly estimated at comparable levels of precision to the C-C and D-D settings. Surprisingly, some parameters in the B-D setting, such as τ_*S*_ and *θ*_*S*_, appear to be even more precisely estimated than in the D-D setting (fig. 5, table S1). A similar observation has been made in the C-A setting in comparison with A-A (Huang *et al*., 2022). There is a slight overestimation of 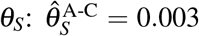 and 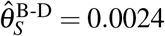 while the true value is 0.002 (table S1).

We obtain a larger estimate of the migration rate when introgression occurs between sister species (B-D) than when it occurs between non-sister species (A-C) even though the introgression probability is the same (φ = 0.2), with 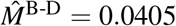 compared with 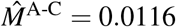 (table S1). Note that in models A and B, introgression occurs at the same time (τ_*X*_ ). The results agree with our analysis of two species that the IIM model (here, model D) recovers more gene flow and provides less biased parameter estimates than the SC model (fig. 3).

#### Both wrong mode and wrong branch (fig. 5, B-C and A-D)

Lastly, we examine the cases when both the mode of gene flow and the lineages involved in gene flow are misspecified (fig. 5: B-C, A-D).

In the B-C setting, introgression occurs from *A* to *T* at τ_*X*_ prior to the divergence of *B* and *C* at τ_*T*_ while the fitting model assumes continuous gene flow from *A* to *B*. The effect of model misspecification is local, affecting mainly τ_*S*_ and *θ*_*S*_. All present-day population sizes (*θ*_*A*_, *θ*_*B*_, *θ*_*C*_ and *θ*_*D*_) as well as τ_*R*_ and *θ*_*R*_ are well estimated. Model C can fit coalescent times during the initial period (0, τ_*T*_ ) well without requiring gene flow. As expected, τ_*T*_ is well estimated and the estimated migration rate is close to zero 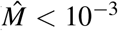 table S1) and is not significant (fig. S7). Small coalescent times *t*_*ab*_ and *t*_*ac*_ due to gene flow in the data generated under model B are then explained by having a more recent divergence time τ_*S*_. We obtain 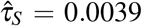, which is much closer to the introgression time τ_*X*_ = 0.003 than the true divergence time τ_*S*_ = 0.006. The root divergence time and population size are slightly affected, with τ_*R*_ slightly overestimated and *θ*_*R*_ slightly underestimated. This pattern is highly similar to the D-C setting discussed above in that failure to detect gene flow has a local effect in the phylogeny, regardless of the mode of gene flow.

In the A-D setting, introgression occurs from *A* to *B*, a non-sister species, while the fitting model assumes continuous gene flow from *A* to its sister lineage *T* . As in the B-C setting, all present-day population sizes, τ_*R*_ and *θ*_*R*_ are well estimated. In the data-generating model, introgression makes some *B* sequences share a more recent common ancestor with those from *A* than from *C* (i.e., *t*_*ab*_ *< t*_*bc*_). We thus expect the divergence time between *B* and *C* be between the introgression time τ_*X*_ and the divergence time τ_*T*_ . We obtain 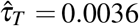, which is between τ_*X*_ = 0.003 and τ_*T*_ = 0.004 (table S1). We find τ_*S*_ to be slightly overestimated, with 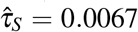 while the true value is 0.006. The biased divergence times are accompanied by a slightly underestimated (table S1). 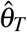 and a slightly overestimated 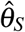(table S1).

In summary, assuming an incorrect model of gene flow and assigning migration to a wrong pair of populations have relatively localized effects on the estimates of divergence time and sizes of populations directly ancestral to gene flow (here, *S*). When gene flow involves ancestral branches in the data-generating model but the fitting model assumes it occurs continuously up to the present, migration rate will be severely underestimated and the Bayesian test may fail to detect gene flow (fig. S7). This pattern corroborates our analysis of the two-species case where the IM and SC models give poorer estimates than the IIM model (fig. 3).

### Misspecified direction of gene flow

Previously, we studied the effects of misspecified direction of gene flow on parameter estimation and Bayesian test of gene flow under the MSC-I model (Thawornwattana *et al*., 2023a). Here, we perform complementary analysis under the MSC-M model. We consider two MSC-M models in figure 5: C (recent gene flow involving non-sister species) and D (ancestral gene flow involving sister species). For each model, there are three variants: inflow (I, *A* → *B*), outflow (O, *B* → *A*), and bidirectional gene flow (B, *A* ⇆ *B*) (fig. 6**a-f**). Results are summarized in figure 6**g&h**. Estimates from the large datasets of *L* = 4,000 loci under models C and D are in tables S2 and S3, respectively.

**Figure 6:**
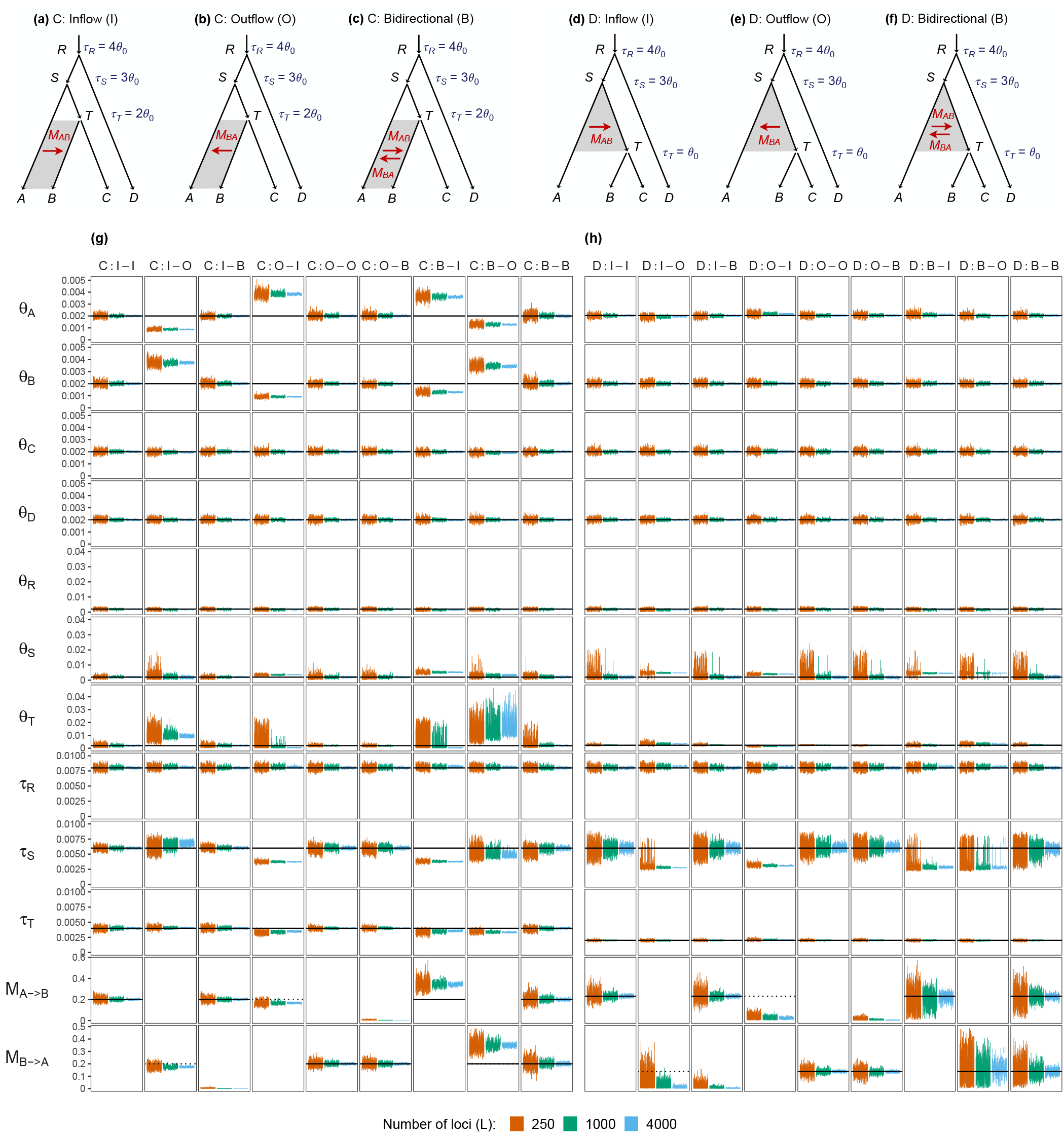
MSC-M models with different directions of gene flow between non-sister lineages *A* and *B* (**a-c**, model C) or between sister lineages *A* and *T* (**d-f**, model D), with either inflow (**b&d**), outflow (**c&e**), or bidirectional gene flow (**d&f**). The notation C:I-O means the data were generated under model C:I (inflow) and analyzed under model C:O (outflow), etc. Parameters used to generate the data are the same as those in figure 5**c&d**. All migration rates were 0.2. For D models, the migration rate *M*_*A*→*T*_ is labelled *M*_*A*→*B*_, etc. for convenience. (**g-h**) The 95% HPD CIs for parameters in 100 replicate datasets of *L* = 250, 1,000, and 4,000 loci for C models (**g**) and D models (**h**). Black solid line indicates the true value. Estimates for the C:I-I and D:I-I settings are identical to those for the C-C and D-D settings, respectively, in figure 5.

#### Analysis under the secondary contact (SC) model (model C, fig. 6g)

In model C, gene flow is recent and between non-sister species *A* and *B*. It may be considered an instance of a secondary contact (SC) model in which gene flow occurs after a period of complete isolation (fig. 1**d**). As a baseline for comparison, we first consider cases where the analysis model is correctly specified, i.e., C:I-I, C:O-O, and C:B-B (fig. 6). In all three cases, all parameters including the migration rates are correctly estimated. The present-day population sizes (*θ*_*A*_, *θ*_*B*_, *θ*_*C*_, *θ*_*D*_) are estimated with narrow CIs while there is considerably uncertainty in the estimates of the ancestral population sizes (*θ*_*T*_, *θ*_*S*_, *θ*_*R*_) (fig. 6**g**). We find that inflow and outflow can be estimated at similar levels of precision, with 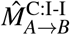 and 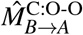 having similar CI widths. It is more difficult to estimate bidirectional gene flow (C:B-B) because there are more parameters in the model. Most parameters have wider CIs, including 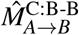 and 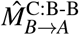.

In the other settings of figure 6**g**, the direction of gene flow is misspecified. Overall, misspecification has local effects on estimation of parameters associated with populations involved in gene flow or their ancestral populations. There is little bias in the estimates of other parameters, such as the present-day population size parameters that are not linked to gene flow (*θ*_*C*_ and *θ*_*D*_), and the root population size and divergence time (τ_*R*_ and *θ*_*R*_) (fig. 6**g**). The divergence time τ_*T*_ of *B* and *C* is also well estimated, likely due to information in *t*_*bc*_ and well-estimated *θ*_*C*_. We focus on parameters that are affected by the misspecified direction of gene flow.

In the C:I-B and C:O-B settings, migration is unidirectional but the analysis model allows for migration in both directions. In those cases, the analysis model is correct but over-parametrized. We recover the correct migration rate in the correct direction while the migration rate in the opposite direction is estimated to be zero (fig. 6**g**, table S2).

The CI widths of all parameters are comparable to those obtained from the unidirectional migration case (compare C:I-B with C:I-I and C:O-B with C:O-O), suggesting that overparameterization has a minor impact on posterior estimates. The cost of including a nonexistent migration rate in the bidirectional model (B) is thus mostly computational.

In the C:I-O and C:O-I settings, migration occurs in one direction but the analysis model assumes the opposite direction. This misspecification of the gene-flow direction should affect estimation of parameters for populations involved in gene flow (*A, B, T* ) and potentially their common ancestor (*S*). When the true recipient population is incorrectly assumed to be a source population, we may expect its population size parameter to be overestimated to account for excess polymorphism. Conversely, a source population incorrectly assumed as a recipient should have its population size parameter underestimated. The results confirm this expectation (fig. 6**g**, table S2). In the C:I-O setting, the recipient population *B* is incorrectly assumed as the source of gene flow. The estimated migration rate 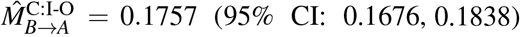 is close to the true migration rate *M*_*A*→*B*_ = 0.2 in the opposite direction, which is significant according to the Bayesian test (fig. S8**a**). The excess polymorphism in *B* due to gene flow is accounted for by having a larger 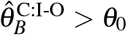 and a smaller 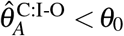. A large 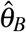 also leads to a grossly overestimated 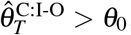, and a slightly overestimated 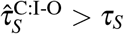.

There are no noticeable biases in other parameters. In the C:O-I setting, the recipient population *A* is incorrectly assumed as the source of gene flow. We obtain a similar pattern in the opposite direction: 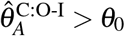 and 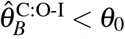, with significant migration detected in the wrong direction (fig. S8**b**). In contrast with the C:I-O setting, the divergence time between *A* and *B* (τ_*S*_) is severely underestimated, with 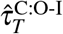 and 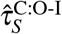 both younger than the true τ_*T*_, and a very short *TS* branch (fig. 6**g**, table S2). These biases in divergence times are associated an underestimation of *θ*_*T*_ and an overestimation of *θ*_*S*_.

In the C:B-I and C:B-O settings, migration occurs in both directions while the analysis model incorrectly assumes unidirectional migration. The results are largely similar to the C:O-I and C:I-O settings, respectively. As expected, the assumed source population (*A* in C:B-I and *B* in C:B-O) has its population size parameter overestimated and the assumed recipient population (*B* in C:B-I and *A* in C:B-O) has its population size parameter underestimated (fig. 6**g**). The bias in 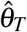 follows that of 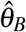. There is a large positive bias in the migration rate estimate, with 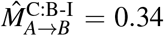 and 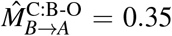 being about twice the estimates from the C:O-I and C:I-O settings 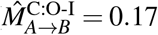 and 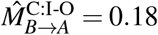 due to a cumulative effect of gene flow in both directions in the data-generating model on reducing the coalescent time *t*_*ab*_ between sequences from *A* and *B*. Consequently, τ_*T*_ and τ_*S*_ are underestimated (table S3).

#### Analysis under the isolation-with-initial-migration (IIM) model (model D, fig. 6h)

Lastly, we consider model D, which has ancestral gene flow between sister lineages *A* and *T* prior to the divergence of *B* and *C* (fig. 6**d-f**). This may be considered as an instance of an IIM model (fig. 1**c**).

When the model is correctly specified (D:I-I, D:O-O, and D:B-B), all parameters are well estimated, with the CIs becoming narrower with the increase of the data size (the number of loci). Again population sizes for ancestral species (*θ*_*S*_, *θ*_*T*_ ) have far wider CIs than those for modern species. Migration rates (*M*_*A*→*B*_ and *M*_*B*→*A*_) as well as τ_*S*_ and *θ*_*S*_ have wider CIs under the D models (fig. 6**h**) than under the corresponding C models (fig. 6**g**). Two factors appear to have made inference of gene flow under model D more challenging than under model C. First, gene flow in D is more ancient. Second, gene flow in D is between sister species.

In the other settings, the direction of gene flow is misspecified (fig. 6**h**). The population sizes for modern species are correctly estimated in all settings (fig. 6**h**), as there is no gene flow during the period (τ_*T*_, 0) in both true and analysis models. The divergence time τ_*T*_ and the population size *θ*_*T*_ are also well estimated in all settings. In the D:I-B and D:O-B settings, assuming bidirectional migration while the true migration is unidirectional has no major impacts on the estimates. The migration rate in the direction that does not occur in the data-generating model is estimated to be zero (fig. 6**h**, table S3).

The D:I-O, D:O-I, D:B-O and D:B-I settings show considerably less biases in parameter estimates than in the corresponding analyses under model C. In particular, τ_*T*_ and *θ*_*T*_ are much better estimated under the D models. Interestingly, when gene flow occurs in one direction but the analysis model only allows gene flow in the opposite direction (D:I-O and D:O-I), the migration rate is estimated to be close to zero as the number of loci increases (fig. 6**h**, table S3) and is often not significant (fig. S8**d-e**). Consequently, the divergence time τ_*S*_ is estimated to be much younger, with 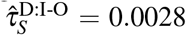 and 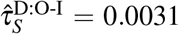 while the true value is τ_*S*_ = 0.006.

No other parameters are affected. This is in contrast to the C:I-O and C:O-I settings where the estimates of migration rate in the wrong direction are positive and close to the true value (fig. 6**g**).

When migration occurs in both directions but the analysis model only allows one direction (D:B-O and D:B-I), we still obtain good estimates of the migration rate in the direction that is allowed, although with a much wider CI compared to that in the D:B-B setting, and the power of the Bayesian test to detect gene flow is relatively moderate in the D:B-O setting fig. S8**f**). The excess polymorphism due to gene flow in the other direction is accounted for by having a much more recent divergence time, with 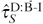 and 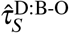 of about 0.003 (cf: the true value of τ_*S*_ = 0.006).

In summary, it is more difficult to detect ancient gene flow between sister species (model D) than recent gene flow between non-sister species (model C). When the direction of gene flow is incorrectly specified, we may still detect significant gene flow in the wrong direction (i.e., a false positive) when gene flow is recent (model C) but not when it is ancestral (model D).

## Discussion

### Summary of key findings

We here summarize five key findings from the current work as well as from previous work on statistical inference under a misspecified gene flow models (Jiao *et al*., 2020; Huang *et al*., 2022; Thawornwattana *et al*., 2023a).

#### Differences between MSC-I and MSC-M models

First, the MSC-I and MSC-M models (e.g., fig. 1) make qualitatively different predictions about how gene flow impacts the distribution of coalescent time between sampled sequences (figs. 4, S2, S4–S6, SI text). In both models, gene flow increases the amount of coalescence between the source and recipient populations and reduces the amount of coalescence within the recipient population, leading to more recent coalescent time. Consider *A* → *B* gene flow. In the MSC-I model, the probability density of the coalescent time *t*_*ab*_ always peaks at the introgression time, with the magnitude depending on the introgression probability (φ), followed by an exponential decay governed by the source population size (e.g., fig. S1). In the MSC-M model, by contrast, the probability density of *t*_*ab*_ has a concave shape with a peak in the middle of the migration period (e.g., fig. S1). As the migration rate (*M*) increases, the peak shifts towards the more recent time and the shape of *t*_*ab*_ becomes more like an exponential decay. At the extreme end of very low gene flow (φ = *M* = 0) or very high gene flow (φ = 1 and *M* → ∞), the two models become essentially equivalent (see SI text). At intermediate levels of gene flow, the mismatches in coalescent time distsibutions lead to biased parameter estimates (figs. 4, S1, S2, S4–S6). Nonetheless, the impact tends to be local in the phylogeny, with most affected parameters being those linked to gene flow, such as the intensity of gene flow (as measured by φ or *M*), population size parameters of the source and recipient populations, as well as their divergence time and ancestral population size. Parameters for other populations on the species phylogeny are usually well estimated.

#### Correspondence between φ and M

Second, we characterize the correspondence between the introgression probability in the MSC-I model (φ) and and the population migration rate in the MSC-M model (*M*), by equating the expected proportion of migrants in the two models: φ = 1 − e^−*w*Δτ^ where 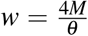 is the mutation-scaled migration rate and Δτ is the time period of migration in the MSC-M model and *θ* is the recipient population size parameter (see Huang *et al*., 2022, eq. 10). This expression provides a reasonable approximation for small values of *M* and φ (figs. 2 and 3). When the MSC-I model is used to analyze the data generated under the MSC-M, the estimated introgression probability φ is always smaller than the expected proportion of migrants (Huang *et al*., 2022). Moreover, a higher amount of gene flow is usually recovered when the data is generated under the SC model than when it is generated under the IM or IIM with the same amount of total gene flow. The estimated introgression time approaches the lower limit of the migration period as the number of sites at each locus approaches infinity. By contrast, when the MSC-M model is used to analyze the data generated under the MSC-I model, the IIM model is able to recover more gene flow across a range of values of φ than the IM and SC models (figs. 2&3 Moreover, the IM and SC models often fail to detect gene flow at extreme values of φ (fig. S3**d**). This appears to be a result of the absence of recent gene flow in the data-generating model. Despite this, species divergence times are often well estimated. In other words, it is easier to infer gene flow under the IIM model than the SC model when the data is generated under the MSC-I model (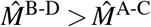; see fig. 5, A-C and B-D), whereas it is easier to infer more recent gene flow under the MSC-I when the data is generated under the SC model than when it is generated under the IIM model (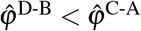; see Huang *et al*., 2022, their fig. 4, C-A and D-B).

#### Incorrect placement of gene flow

Third, when gene flow is assigned to a wrong pair of populations in the phylogeny but the mode of gene flow is correctly assumed, estimates of the migration rate and divergence/introgression times can be highly biased. When gene flow is recent and between non-sister species but the fitting model assumes ancestral gene flow between sister lineages (e.g. fig. 5, C-D), we may still be able to detect gene flow. Furthermore, in the MSC-M model, the species divergence times and the migration period may be severely underestimated, with biased estimates of the source and recipient population sizes. In the MSC-I model, the introgression time tends to collapse onto the species divergence time (Huang *et al*., 2022, their fig. 4, A-B).

When gene flow is ancestral and between sister lineages but the fitting model assumes recent gene flow between non-sister species (e.g. fig. 5, D-C), we may not detect any gene flow, and consequently, the species divergence time is underestimated. Moreover, in the MSC-I model, the estimated introgression time tends to get stuck at the species divergence time (Huang *et al*., 2022, their fig. 4, B-A).

#### Misspecified direction of gene flow

Fourth, when the direction of gene flow is misspecified, recent gene flow between non-sister species can still be detected despite the misspecification of direction (e.g. fig. 6, C:I-O, C:O-I; also see Thawornwattana *et al*., 2023a, fig. S4, I-O and O-I), but the estimates of species divergence times and effective population sizes for the source and recipient populations are biased. By contrast, when gene flow is ancestral and between sister species, we do not detect gene flow in the opposite direction (fig. S8). This is partly because parameters associated with modern populations are well estimated (e.g. fig. 6, D:I-O, D:O-I), and the excess polymorphism in the recipient population is accounted for by underestimating the species divergence times.

#### Bayesian test for detecting gene flow

Lastly, the Bayes factor can be used to test whether the estimated gene flow is significantly larger than zero. For nested models of gene flow, the Bayes factor can be estimated using the Savage–Dickey density ratio (Dickey, 1971) calculated from posterior MCMC samples (Ji *et al*., 2023). We find that the IM and SC models may not detect gene flow when used to analyze data generated under the MSC-I model while the IIM model always does (fig. S3). Moreover, we still detect continuous migration in the wrong direction when gene flow is recent and between non-sister species (fig. S8: C:I-O and C:O-I) but not when gene flow is ancestral and between sister species (fig. S8: D:I-O and D:O-I). Similarly, we can still detect pulse introgression when the direction of introgression is misspecified under the MSC-I model (Thawornwattana *et al*., 2023a).

### Recommendations for the use of models of gene flow in analysis of genomic data

The MSC-I and MSC-M are two extreme modes of gene flow models. In reality, we expect a variety of intermediate modes by which gene flow might occur between species or populations. For example, the intensity of gene flow may vary over time as species expand and move across space, impacting their opportunities to meet and hybridize. What we observe in genomic sequence data is an outcome of long-term interactions among multiple factors and processes including natural selection purging introgressed alleles, recombination breaking the linkage to loci under selection, and genetic drift (Martin and Jiggins, 2017; Moran *et al*., 2021). Thus the amount of gene flow estimated from the genomic data under the MSC-M (migration rate per generation, *M*) and MSC-I (introgression probability, φ) models reflects ‘effective’ gene flow, and is different from, say, the proportion of F1 hybrids in the contact zone. Which of the two modes of gene flow is more suitable will depend on the system under study. The MSC-I model should be more suitable for ancient gene flow or gene flow that occurred over a brief time period, while the MSC-M model is more suitable for on-going gene flow between extant populations or gene flow that occurs over an extended time period. In this regard, a few thousand generations may be represented as a pulse on an evolutionary time scale. Our results in this study (see also Huang *et al*., 2022; Thawornwattana *et al*., 2023a) suggest that the MSC-I model is more robust to various kinds of model misspecifications, and should in general be recommended. The MSC-M model may be useful if there is substantive evidence for continuous gene flow over extended time periods.

Bayesian model selection based on Bayes factor can be used to test whether the detected gene flow is statistically significant (see ‘*Bayesian test for detecting gene flow*’). An application of such an approach to the genomic data from the *Anopheles gambiae* complex of African mosquitoes suggested that the MSC-I model fits the data better than the MSC-M model (Flouri *et al*., 2023). But regardless of whether the mode of gene flow is correctly specified, this and previous studies (Huang *et al*., 2022; Thawornwattana *et al*., 2023a) have found that parameters for lineages not involved in gene flow, such as the root age and root population size, tend to be well estimated under either model. We also recommend the use of the bidirectional gene-flow model when the direction of gene flow is not known *a priori*.

Summary methods may be considered approximations to full-likelihood methods under the multispecies coalescent without making explicit assumptions about the underlying processes that generate gene trees and sequence data. For instance, the *D* statistic (Green *et al*., 2010; Durand *et al*., 2011), the *f* statistic (Reich *et al*., 2009; Patterson *et al*., 2012), and HyDe (Blischak *et al*., 2018; Kubatko and Chifman, 2019) use site pattern counts pooled across the genome or from a large genomic region. These patterns provide information on relative frequencies of gene trees under a variety of mutation models. While there is symmetry in the expected frequencies of discordant gene trees if ancestral polymorphism is the only factor causing genealogical fluctuations across the genome, gene flow between species breaks this symmetry, and this fact is used to detect gene flow. Other summary methods make use of gene-tree frequencies directly, including the Δ statistic (Huson *et al*., 2005), SNAQ (Solis-Lemus and Ane, 2016) and PHRAPL (Jackson *et al*., 2017). For both site pattern-based and gene tree-based methods, summarizing sequence alignments from multiple samples and multiple species at thousands of genomic loci into two or three proportions (of site patterns or of gene trees) is dramatic and leads to a considerable information loss. Most summary methods cannot detect gene flow between sister lineages, and cannot infer the direction, timing or amount of gene flow, or estimate key parameters such as effective population sizes and species divergence times (Jiao *et al*., 2021; Huang *et al*., 2022).

Summary methods, however, do have a computational advantage over full-likelihood methods and can be easily applied to genomic data to infer broad patterns of gene flow in a large phylogeny. Nonetheless, we suggest that likelihood implementations of the MSC-I and MSC-M models may play an important role that compliments and extends summary methods. Full-likelihood methods represent highly parametric models, based on modelling the biological process of reproduction, species divergence and gene flow, and provide a powerful framework for characterizing the rich and complex of species divergence and gene flow using genomic data (Jiao *et al*., 2021).

## Materials and Methods

### Two species case: Asymptotic analysis

The case of two species when the data consist of one sequence per species is simple enough to yield analytical solutions. We consider estimation of parameters under the three MSC-M models of figure 1**b-d** when data of an infinite number of loci (*L* → ∞) are generated under the MSC-I model of figure 1**a**. We obtained the limit of the MLEs, 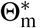, by minimizing the KL divergence (eq. 1). As *L* → ∞, the data are represented by the distribution of the number of differences between two sequences at the *n* sites, and maximizing the likelihood is equivalent to minimizing the KL divergence. Optimization was achieved using a C program that implements the BFGS algorithm from PAML (Yang, 2007) and was previously used by Huang *et al*. (2022). The program is available at https://github.com/ythaworn/iimmsci2s. For each model and each value of φ, we ran the optimization multiple times and used the run with the lowest KL value. We excluded runs with parameter values on the optimization boundaries.

### Two species case: Simulation

We used simulation to verify and extend our asymptotic analysis. We simulated multilocus sequence data under the MSC-I model of figure 1**a** and analyzed them under the IM, IIM and SC models (figure 1**b-d**).

We used two values of population sizes on the species tree: *θ*_*A*_ = *θ*_*X*_ = *θ*_*R*_ = *θ*_0_ = 0.002 (thin branches) and *θ*_*B*_ = *θ*_*Y*_ = *θ*_1_ = 0.01 (thick branches). Introgression occurred from species *A* to *B* with probability φ = 0.2 at time τ_*X*_ = *θ*_0_ after species divergence at time τ_*R*_ = 2*θ*_0_. In the base case, we assumed φ = 0.2 and the data consisted of *L* = 4,000 loci, with *S* = 4 sequences per species, and *n* = 1,000 sites per sequence. We varied the following four factors one at a time, keeping other parameters fixed: the number of sites per sequence (*n*), the number of sequences per species (*S*), the number of loci (*L*), and the introgression probability (φ).

The values used were *n* =250, 1,000, 4,000, 16,000, 64,000; *S* = 1, 2, 4, 8, 16; *L* =250, 500, 1,000, 2,000, 4,000, 8,000; and φ =0.01, 0.05, 0.1, 0.2, 0.3, 0.4, 0.5, and 0.7. For each setting, we simulated 30 replicate datasets. With those four factors (*n, S, L, M*), and 30 replicates each, there were (5 + 5 + 6 + 8 − 3) *×* 30 = 630 datasets in total. The subtraction by 3 accounted for the fact that all four factors shared the same base case (*n* = 1,000, *S* = 4, *L* = 4,000, φ = 0.2). To generate sequence data, we first generated a gene tree with coalescent times at each locus and then simulated sequences along branches of the gene tree under the JC model (Jukes and Cantor, 1969). Sequences at the tips of the tree became data at the locus. Simulation was done using the `simulate` option in BPP v4.7.0 (Flouri *et al*., 2018, 2020, 2023).

Each dataset was analyzed under the IM, IIM, and SC models of figure 1**b-d** to estimate parameters using BPP v4.7.0 (Flouri *et al*., 2023). The JC mutation model was assumed. We assigned gamma priors to population size parameters (*θ* ), the root age (τ_*R*_) on the species tree and the migration rate: *θ* ∼ *G*(2, 200) with mean 2*/*200 = 0.01, τ_*R*_ ∼ *G*(4, 200) with mean 4*/*200 = 0.02, and *M* ∼ *G*(2, 10) with mean 2*/*10 = 0.2. For each fitting model, we perform two independent runs of MCMC, each with 32,000 iterations of burnin and 10^6^ iterations of the main chain. Samples were recorded every 100 iterations. With three fitting models and two MCMC runs per dataset, there were 3 *×* 2 *×* 630 = 3,780 MCMC runs in total. Each run of the base case took about 80 hours (hrs) and 2G of memory while the most expensive runs (*L* = 8,000 or *S* = 8) took about 200 hr and 4G of memory. For datasets with *S* = 16, we allowed the MCMC run for up to 300 hrs and 8G of memory, which was about 4 *×* 10^5^ iterations.

### Four species case: Wrong migration branch and wrong mode of gene flow

Data were simulated under the four models (A-D) in a phylogeny for four species in figure 5 and analyzed under models C and D. Models A and B assumes pulse introgression while model C and D assumes continuous migration. Gene flow occurred either between non-sister species (models A and C) or between sister species (models B and D). Parameters are shown in figure 5. All population sizes were assumed to be *θ*_0_ = 0.002. For models A and B, we used the introgression probability φ = 0.2. For models C and D, we assumed the migration rate *M* = 0.2 migrants per generation. Each dataset consisted of *S* = 4 sequences per species per locus, each of length *n* = 500 sites. We varied the number of loci: *L* =250, 1,000, and 4,000. For each setting, we simulated 100 replicate datasets.

Each dataset was analyzed under both models C and D (fig. 5**c&d**) using BPP v4.7.0 (Flouri *et al*., 2023). There are eight settings in total: A-C, B-C, C-C, D-C, A-D, B-D, C-D, and D-D. Settings C-C and D-D serve a reference when the model is correctly specified. In settings C-D and D-C, the population pair (or branches in the phylogeny) involved in migration is misspecified. In settings A-C and B-D, gene flow occurs as pulse introgression but is misspecified as continuous migration. Finally, settings B-C and A-D have a combined feature of an incorrect mode of gene flow assigned to a wrong branch. Gamma priors were assigned to population sizes, root age and migration rate as *θ* ∼ *G*(2, 200) with mean 0.01, τ_*R*_ ∼ *G*(4, 200) with mean 0.02, and *M* ∼ *G*(2, 10) with mean 0.2. With four data-generating models, three values of *L* and 100 replicates, there were 4 *×* 3 *×* 100 = 1,200 datasets in total. We performed two independent runs of MCMC, each with 10,000 iterations of burnin and 10^6^ iterations of the main chain. Samples were recorded every 100 iterations. With two fitting models (C and D) for each dataset, there were 2 *×* 2 *×* 1,200 = 4,800 MCMC runs in total. The running time was about 20-30 hrs for datasets with *L* = 250, 80 hrs for *L* = 1,000, and 260 hrs for *L* = 4,000.

### Simulation in the four species case: Misspecified direction of gene flow

To study the effect of incorrectly assumed direction of gene flow under the MSC-M model, we simulated sequence data using models C and D of figure 5, but with three specifications concerning the direction of gene flow: inflow (I, *A* → *B*), outflow (O, *B* → *A*), and bidirectional gene flow (B, *A* ⇆ *B*) (figure 6**a-f**). We used the same parameter values as in the previous section (fig. 5**c-d**)). Each simulated dataset is analyzed assuming the three variants of the MSC-I model (I, O, and B), generating nine settings for model C (e.g., C:I-O) and nine settings for model D (e.g., D:I-O).

We used three values of for the number of loci: *L* = 250, 1,000, 4,000. For model C, with 3 data-generating models, three values of *L* and 100 replicates, there were 3 *×* 3 *×* 100 = 900 datasets in total. MCMC setup was the same as before. With three fitting models (I, O, and B), two independent MCMC runs per dataset, the total number of MCMC runs was 3 *×* 2 *×* 900 = 5,400. Similarly for model D, there were 900 datasets and 5,400 MCMC runs in total. We reused the datasets for C:I and D:I from the previous section.

### Bayesian test of gene flow

We conducted the Bayesian test of gene flow (Ji *et al*., 2023) to assess whether there is significant evidence in the data for gene flow (Thawornwattana *et al*., 2023a). For example, for unidirectional gene flow (figs. 1, 5), the null hypothesis of no gene flow, *H*_0_ : *M*_*A*→*B*_ = 0 may be compared with the alternative hypothesis of gene flow, *H*_1_ : *M*_*A*→*B*_ *>* 0, via the Bayes factor *B*_10_. As the two hypotheses are nested, *B*_10_ may be approximated by the Savage-Dickey density ratio, approximately given as 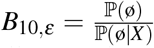, where ℙ (ø) is the probability for the null interval, ø : 0 *< M*_*A*→*B*_ *< ε*, under the prior distribution, and ℙ (ø|*X* ) is the corresponding posterior probability. The null interval is part of the parameter space for *H*_1_ that represents the null hypothesis. We used *ε* = 0.01 and 0.001. *B*_10,*ε*_ was calculated by processing a posterior MCMC sample under *H*_1_ (Ji *et al*., 2023). *B*_10_ *>* 100 is considered strong evidence in favor of *H*_1_, similar to the 1% significance level in hypothesis testing. The power of the test is defined as the proportion of replicate datasets in which *B*_10_ *>* 100.

## Acknowledgments

This studyhasbeensupported by Biotechnology and Biological Sciences Research Council (BBSRC) grants (BB/T003502/1, BB/X007553/1) and a Natural Environment Research Council (NERC) grant (NE/X002071/1) to Z.Y., as well as by Harvard University.

## Data Availability

Simulated datasets are available in Zenodo at https://doi.org/10.5281/zenodo.11182437.

