## supplemental information for "Inference of continuous gene flow between species under misspecified models"

### SI text: Coalescent time distributions under the MSC-I and MSC-M models for two species

We derive probability densities of coalescent times between two sequences under the MSC-I and MSC-M models for two species of figure 1**a**-**d**. Our asymptotic analysis is based on $f(t_{ab})$, the distribution of coalescent time for two sequences sampled from the two species. We also derive $f(t_{aa})$ and $f(t_{bb})$ for two sequences sampled from the same species, as comparison of the true and fitting distributions of those coalescent times provides important insights into Bayesian parameter estimation under misspecified models (e.g. fig. 4).

#### *Density of coalescent time under the MSC-I model of figure 1a*

In the MSC-I model of figure 1**a**, introgression occurs from species $A$ into $B$ at time $\tau_X$ with probability $\varphi$. With at least two sequences sampled per species, there are eight parameters in total: $\Theta_{\rm i} = \{\varphi, \tau_X, \tau_R, \theta_A, \theta_B, \theta_X, \theta_Y, \theta_R\}$. However, some parameters may be unidentifiable if only one sequence is sampled per species (Yang and Flouri, 2022).

The coalescent time between one sequence from $A$ and one from $B$ has density (e.g., Jiao *et al.*, 2020)

$$f_{{\rm i},ab}(t) = \begin{cases} \varphi \frac{2}{\theta_X} \mathrm{e}^{-\frac{2}{\theta_X}(t-\tau_X)}, & \text{if } \tau_X < t \le \tau_R, \\ \left[\varphi\, \mathrm{e}^{-\frac{2}{\theta_X}(\tau_R-\tau_X)} + (1-\varphi)\right] \frac{2}{\theta_R} \mathrm{e}^{-\frac{2}{\theta_R}(t-\tau_R)}, & \text{if } t > \tau_R. \end{cases} \tag{S1}$$

This is independent of $\theta_A, \theta_B$ and $\theta_Y$.

For two sequences from $A$, the density of coalescent time is piecewise exponential

$$f_{{\rm i},aa}(t) = \begin{cases} \frac{2}{\theta_A} \mathrm{e}^{-\frac{2}{\theta_A}t}, & \text{if } 0 < t \le \tau_X, \\ \mathrm{e}^{-\frac{2}{\theta_A}\tau_X} \frac{2}{\theta_X} \mathrm{e}^{-\frac{2}{\theta_X}(t-\tau_X)}, & \text{if } \tau_X < t \le \tau_R, \\ \mathrm{e}^{-\frac{2}{\theta_A}\tau_X} \mathrm{e}^{-\frac{2}{\theta_X}(\tau_R-\tau_X)} \frac{2}{\theta_R} \mathrm{e}^{-\frac{2}{\theta_R}(t-\tau_R)}, & \text{if } t > \tau_R. \end{cases} \tag{S2}$$

This is independent of $\theta_B, \theta_Y$ and $\varphi$.

Lastly, for two sequences from $B$, we have

$$f_{{\rm i},bb}(t) = \begin{cases} \frac{2}{\theta_B} \mathrm{e}^{-\frac{2}{\theta_B}t}, & \text{if } 0 < t \le \tau_X, \\ \mathrm{e}^{-\frac{2}{\theta_B}\tau_X} \left[(1-\varphi)^2 \frac{2}{\theta_Y} \mathrm{e}^{-\frac{2}{\theta_Y}(t-\tau_X)} + \varphi^2 \frac{2}{\theta_X} \mathrm{e}^{-\frac{2}{\theta_X}(t-\tau_X)}\right], & \text{if } \tau_X < t \le \tau_R, \\ \mathrm{e}^{-\frac{2}{\theta_B}\tau_X} \left[(1-\varphi)^2 \mathrm{e}^{-\frac{2}{\theta_Y}(\tau_R-\tau_X)} + \varphi^2 \mathrm{e}^{-\frac{2}{\theta_X}(\tau_R-\tau_X)} + 2\varphi(1-\varphi)\right] \frac{2}{\theta_R} \mathrm{e}^{-\frac{2}{\theta_R}(t-\tau_R)}, & \text{if } t > \tau_R. \end{cases} \tag{S3}$$

This is independent of $\theta_A$.

In this work, we always assume $\theta_A = \theta_X$ and $\theta_B = \theta_Y$, i.e., there is no change in the population size after introgression.

#### *Density of coalescent time between two sequences under the MSC-M models (IM, IIM and SC) of figure 1b-d*

In the IM model (fig. 1**b**), the two species ($A$ and $B$) diverge at time $\tau_R$ and there is continuous gene flow from $A$ to $B$ at rate of $M_{A\to B}$ migrants per generation since $\tau_R$. In the IIM model (fig. 1**c**) gene flow continues to occur from $A$ to $B$ after species divergence but stops at time $\tau_T > 0$ (Costa and Wilkinson-Herbots, 2017). In the SC model, (fig. 1**d**), the two species are initially completely isolated after their divergence but gene flow starts to occur from $A$ to $B$ at rate $M_{A\to B}$ from time $\tau_T > 0$ to the present (Costa and Wilkinson-Herbots, 2021). The IM model (fig. 1**b**) can be viewed as a special case of the IIM model (fig. 1**c**) with $\tau_T = 0$ or as a special case of the SC model (fig. 1**d**) with $\tau_T = \tau_R$. Here we derive the densities for the coalescent time between two sequences ($t_{aa}, t_{ab}, t_{bb}$) under the IIM and SC models.

Consider the IIM model of figure 1**b**. The parameter vector is $\Theta_{\rm IIM} = \{M, \tau_T, \tau_R, \theta_A, \theta_B, \theta_T, \theta_R\}$. The backwards-in-time process of coalescent and migration in time interval $(\tau_T, \tau_R)$ for two sequences is described by a Markov chain with four states: $AB$, $AA$, $BB$ and $A$ (Notohara, 1990). Here $AA$ means two sequences in $A$ and $A$ means one sequence in $A$ (i.e., a coalescent event has occurred, reducing the number of sequences from two to one). Note that the $AB$-to-$AA$ transition means migration of a sequence from $A$ to $B$ in the real world (forward time). The rate matrix of this Markov chain is

$$
Q = \begin{array}{c|cccc}
 & AA & AB & BB & A \\
\hline
AA & -\frac{2}{\theta_A} & 0 & 0 & \frac{2}{\theta_A} \\
AB & w & -w & 0 & 0 \\
BB & 0 & 2w & -2w-\frac{2}{\theta_B} & \frac{2}{\theta_B} \\
A & 0 & 0 & 0 & 0
\end{array} \tag{S4}
$$

where $w = m_{A\to B}/\mu = 4M_{A\to B}/\theta_B$ is the mutation-scaled migration rate; see, e.g., Herbots (1997), Wilkinson-Herbots (2008), Hobolth *et al.* (2011), and Jiao *et al.* (2020). Coalescence occurs at rate $\frac{2}{\theta_A}$ in $A$ and $\frac{2}{\theta_B}$ in $B$. Time is measured in the expected number of mutations per site. The matrix $Q$ has eigenvalues $\lambda_1 = 0$, $\lambda_2 = -\frac{2}{\theta_A}$, $\lambda_3 = -w$, and $\lambda_4 = -\frac{2}{\theta_B} - 2w$. Let the transition probability matrix over time $t$ be $P(t) = \{p_{ij}(t)\} = \mathrm{e}^{Qt}$, where $p_{ij}(t)$ is the probability that the Markov chain will be in state $j$ time $t$ later given that it is in state $i$ at time 0. This is

$$
P(t) = \begin{bmatrix}
\mathrm{e}^{-\frac{2}{\theta_A}t} & 0 & 0 & 1-\mathrm{e}^{-\frac{2}{\theta_A}t} \\
\frac{\theta_A w}{2-\theta_A w}\left(\mathrm{e}^{-wt}-\mathrm{e}^{-\frac{2}{\theta_A}t}\right) & \mathrm{e}^{-wt} & 0 & 1-\frac{2\,\mathrm{e}^{-wt}-\theta_A w\,\mathrm{e}^{-\frac{2}{\theta_A}t}}{2-\theta_A w} \\
h(t) & \frac{2\theta_B w}{2+\theta_B w}\left(\mathrm{e}^{-wt}-\mathrm{e}^{-\frac{2}{\theta_B+2w}t}\right) & \mathrm{e}^{-\frac{2}{\theta_B+2w}t} & g(t) \\
0 & 0 & 0 & 1
\end{bmatrix}, \tag{S5}
$$

where

$$
h(t) = \frac{w^2\theta_A\theta_B}{(2-w\theta_A)(2+w\theta_B)(\theta_A-\theta_B+w\theta_A\theta_B)}
\left[(2-w\theta_A)\theta_B e^{-(\frac{2}{\theta_B}+2w)t} - \theta_A(2+w\theta_B)e^{-\frac{2}{\theta_A}t} + 2(\theta_A-\theta_B+w\theta_A\theta_B)e^{-wt}\right] \tag{S6}
$$

and

$$
g(t) = 1 - h(t) - \frac{2\theta_B w}{2+\theta_B w}\left(\mathrm{e}^{-wt}-\mathrm{e}^{-\frac{2}{\theta_B+2w}t}\right) - \mathrm{e}^{-\frac{2}{\theta_B+2w}t}. \tag{S7}
$$

Let $\tilde{P}(t)$ denote the transition probability matrix during $(\tau_T, \tau_R)$. This is the same as $P(t)$ except that population $T$ is the recipient of gene flow instead of $B$, with $\theta_B$ replaced by $\theta_T$. Under the IIM model (fig. 1**c**), the probability density of the coalescent time $t_{ab}$ is

$$
\begin{aligned}
f_{\text{IIM},ab}(t) &= \begin{cases} \tilde{P}_{AT,AA}(t-\tau_T)\frac{2}{\theta_A}, & \text{if } \tau_T < t \le \tau_R, \\ [1-\tilde{P}_{AT,A}(\tau_R-\tau_T)]\frac{2}{\theta_R}\,\mathrm{e}^{-\frac{2}{\theta_R}(t-\tau_R)}, & \text{if } t > \tau_R \end{cases} \\
&= \begin{cases} \frac{2w}{2-\theta_A w}\left[\mathrm{e}^{-w(t-\tau_T)} - \mathrm{e}^{-\frac{2}{\theta_A}(t-\tau_T)}\right], & \text{if } \tau_T < t \le \tau_R, \\ \left[\frac{2}{2-\theta_A w}\,\mathrm{e}^{-w(\tau_R-\tau_T)} - \frac{\theta_A w}{2-\theta_A w}\,\mathrm{e}^{-\frac{2}{\theta_A}(\tau_R-\tau_T)}\right]\frac{2}{\theta_R}\,\mathrm{e}^{-\frac{2}{\theta_R}(t-\tau_R)}, & \text{if } t > \tau_R. \end{cases}
\end{aligned} \tag{S8}
$$

This is independent of $\theta_B$. Note that it is a function of $w = 4M_{AT}/\theta_T$ but not of $M_{AT}$ and $\theta_T$ individually.

The other two coalescent times, $t_{aa}$ and $t_{bb}$, have densities

$$
\begin{aligned}
f_{\text{IIM},aa}(t) &= \begin{cases} \frac{2}{\theta_A}\,\mathrm{e}^{-\frac{2}{\theta_A}t}, & \text{if } 0 < t \le \tau_T, \\ P_{AA,AA}(\tau_T)\left[\tilde{P}_{AA,AA}(t-\tau_T)\frac{2}{\theta_A} + \tilde{P}_{AA,AT}(t-\tau_T)\frac{2}{\theta_T}\right], & \text{if } \tau_T < t \le \tau_R, \\ P_{AA,AA}(\tau_T)\left[\tilde{P}_{AA,AA}(\tau_R-\tau_T) + \tilde{P}_{AA,AT}(\tau_R-\tau_T)\right]\frac{2}{\theta_R}\,\mathrm{e}^{-\frac{2}{\theta_R}(t-\tau_R)}, & \text{if } t > \tau_R \end{cases} \\
&= \begin{cases} \frac{2}{\theta_A}\,\mathrm{e}^{-\frac{2}{\theta_A}t}, & \text{if } 0 < t \le \tau_R, \\ \mathrm{e}^{-\frac{2}{\theta_A}(\tau_R-\tau_T)}\,\frac{2}{\theta_R}\,\mathrm{e}^{-\frac{2}{\theta_R}(t-\tau_R)}, & \text{if } t > \tau_R, \end{cases}
\end{aligned} \tag{S9}
$$

which depends on $\theta_A$, $\theta_R$, and $\tau_R$ only, and

$$
f_{\text{IIM},bb}(t) = \begin{cases} \frac{2}{\theta_B}\,\mathrm{e}^{-\frac{2}{\theta_B}t}, & \text{if } 0 < t \le \tau_T, \\ \mathrm{e}^{-\frac{2}{\theta_B}\tau_T}\left[\tilde{P}_{TT,AA}(t-\tau_T)\frac{2}{\theta_A} + \tilde{P}_{TT,AT}(t-\tau_T)\frac{2}{\theta_T}\right], & \text{if } \tau_T < t \le \tau_R, \\ \mathrm{e}^{-\frac{2}{\theta_B}\tau_T}\left[1-\tilde{P}_{TT,A}(\tau_R-\tau_T)\right]\frac{2}{\theta_R}\,\mathrm{e}^{-\frac{2}{\theta_R}(t-\tau_R)}, & \text{if } t > \tau_R \end{cases}
$$

$$
= \begin{cases} \frac{2}{\theta_B}\,\mathrm{e}^{-\frac{2}{\theta_B}t}, & \text{if } 0 < t \le \tau_T, \\ \mathrm{e}^{-\frac{2}{\theta_B}\tau_T}\left[\tilde{h}(t-\tau_T)\frac{2}{\theta_A} + \mathrm{e}^{-(\frac{2}{\theta_T}+2w)(t-\tau_T)}\,\frac{2}{\theta_T}\right], & \text{if } \tau_T < t \le \tau_R, \\ \mathrm{e}^{-\frac{2}{\theta_B}\tau_T}\left[1-\tilde{g}(\tau_R-\tau_T)\right]\frac{2}{\theta_R}\,\mathrm{e}^{-\frac{2}{\theta_R}(t-\tau_R)}, & \text{if } t > \tau_R, \end{cases} \tag{S10}
$$

which depends on all seven parameters in $\Theta_{\text{IIM}}$.

As $M \to \infty$, the IIM model becomes equivalent to the MSC-I model (fig. 1**a**) with $\varphi = 1$ provided $\tau_T = \tau_X$ and $\theta_T = \theta_X$. Also the IIM model with $M = 0$ and any value of $\tau_T$ is equivalent to the MSC-I model with $\varphi = 0$.

For the IM model (fig. 1**b**), the densities $f_{\text{IM},ab}(t)$, $f_{\text{IM},aa}(t)$ and $f_{\text{IM},bb}(t)$ are given by setting $\tau_T = 0$ and $w = 4M_{AB}/\theta_B$ in $f_{\text{IIM},ab}(t)$, $f_{\text{IIM},aa}(t)$ and $f_{\text{IIM},bb}(t)$, respectively.

Finally, under the SC model (fig. 1**c**), the parameter vector is $\Theta_{\text{SC}} = \{M, \tau_T, \tau_R, \theta_A, \theta_B, \theta_T, \theta_R\}$. The coalescent-with-migration process over the time interval (0, $\tau_T$) is described by the Markov chain of eq. S4. The probability density of the coalescent time $t_{ab}$ is

$$
f_{\text{SC},ab}(t) = \begin{cases} P_{AB,AA}(t)\frac{2}{\theta_A}, & \text{if } 0 < t < \tau_T, \\ P_{AB,AA}(\tau_T)\frac{2}{\theta_A}\,\mathrm{e}^{-\frac{2}{\theta_A}(t-\tau_T)}, & \text{if } \tau_T < t < \tau_R, \\ \left[P_{AB,AA}(\tau_T)\,\mathrm{e}^{-\frac{2}{\theta_A}(\tau_R-\tau_T)} + P_{AB,AB}(\tau_T)\right] \times \frac{2}{\theta_R}\,\mathrm{e}^{-\frac{2}{\theta_R}(t-\tau_R)}, & \text{if } t > \tau_R \end{cases}
$$

$$
= \begin{cases} \frac{w\theta_A}{2-w\theta_A}\left[\mathrm{e}^{-wt} - \mathrm{e}^{-\frac{2}{\theta_A}t}\right]\frac{2}{\theta_A}, & \text{if } 0 < t < \tau_T, \\ \frac{w\theta_A}{2-w\theta_A}\left[\mathrm{e}^{-w\tau_T} - \mathrm{e}^{-\frac{2}{\theta_A}\tau_T}\right]\frac{2}{\theta_A}\,\mathrm{e}^{-\frac{2}{\theta_A}(t-\tau_T)}, & \text{if } \tau_T < t < \tau_R, \\ \left[\frac{w\theta_A}{2-w\theta_A}\left[\mathrm{e}^{-w\tau_T} - \mathrm{e}^{-\frac{2}{\theta_A}\tau_T}\right]\mathrm{e}^{-\frac{2}{\theta_A}(\tau_R-\tau_T)} + \mathrm{e}^{-w\tau_T}\right] \times \frac{2}{\theta_R}\,\mathrm{e}^{-\frac{2}{\theta_R}(t-\tau_R)}, & \text{if } t > \tau_R, \end{cases} \tag{S11}
$$

which is independent of $\theta_B$ and $\theta_T$. The coalescent time $t_{aa}$ has density

$$
f_{\text{SC},aa}(t) = \begin{cases} P_{AA,AA}(t)\frac{2}{\theta_A}, & \text{if } 0 < t < \tau_T, \\ P_{AA,AA}(\tau_T)\frac{2}{\theta_A}\,\mathrm{e}^{-\frac{2}{\theta_A}(t-\tau_T)}, & \text{if } \tau_T < t < \tau_R, \\ \left[P_{AA,AA}(\tau_T)\,\mathrm{e}^{-\frac{2}{\theta_A}(\tau_R-\tau_T)}\right] \times \frac{2}{\theta_R}\,\mathrm{e}^{-\frac{2}{\theta_R}(t-\tau_R)}, & \text{if } t > \tau_R \end{cases}
$$

$$
= \begin{cases} \frac{2}{\theta_A}\,\mathrm{e}^{-\frac{2}{\theta_A}t}, & \text{if } 0 < t \le \tau_R, \\ \mathrm{e}^{-\frac{2}{\theta_A}(\tau_R-\tau_T)}\,\frac{2}{\theta_R}\,\mathrm{e}^{-\frac{2}{\theta_R}(t-\tau_R)}, & \text{if } t > \tau_R, \end{cases} \tag{S12}
$$

which is independent of $\theta_B$ and $\theta_T$. The density is identical to $f_{\text{IIM},aa}(t)$.

Lastly, the coalescent time $t_{bb}$ has density

$$
f_{\text{SC},ab}(t) = \begin{cases} P_{BB,AA}(t)\frac{2}{\theta_A} + P_{BB,BB}(t)\frac{2}{\theta_B}, & \text{if } 0 < t < \tau_T, \\ P_{BB,AA}(\tau_T)\frac{2}{\theta_A}\,\mathrm{e}^{-\frac{2}{\theta_A}(t-\tau_T)} + P_{BB,BB}(\tau_T)\frac{2}{\theta_T}\,\mathrm{e}^{-\frac{2}{\theta_T}(t-\tau_T)}, & \text{if } \tau_T < t < \tau_R, \\ \left[P_{BB,AA}(\tau_T)\,\mathrm{e}^{-\frac{2}{\theta_A}(\tau_R-\tau_T)} + P_{BB,AB}(\tau_T) + P_{BB,BB}(\tau_T)\,\mathrm{e}^{-\frac{2}{\theta_T}(\tau_R-\tau_T)}\right] \times \frac{2}{\theta_R}\,\mathrm{e}^{-\frac{2}{\theta_R}(t-\tau_R)}, & \text{if } t > \tau_R \end{cases}
$$

$$
= \begin{cases} h(t)\frac{2}{\theta_A} + \mathrm{e}^{-(\frac{2}{\theta_B}+2w)t}\,\frac{2}{\theta_B}, & \text{if } 0 < t < \tau_T, \\ h(\tau_T)\frac{2}{\theta_A}\,\mathrm{e}^{-\frac{2}{\theta_A}(t-\tau_T)} + \mathrm{e}^{-(\frac{2}{\theta_B}+2w)\tau_T}\,\frac{2}{\theta_T}\,\mathrm{e}^{-\frac{2}{\theta_T}(t-\tau_T)}, & \text{if } \tau_T < t < \tau_R, \\ \left[h(\tau_T)\,\mathrm{e}^{-\frac{2}{\theta_A}(\tau_R-\tau_T)} + \frac{2w\theta_B}{2+w\theta_B}\left\{\mathrm{e}^{-w\tau_T} - \mathrm{e}^{-(\frac{2}{\theta_B}+2w)\tau_T}\right\} + \mathrm{e}^{-(\frac{2}{\theta_B}+2w)\tau_T}\,\mathrm{e}^{-\frac{2}{\theta_T}(\tau_R-\tau_T)}\right] & \\ \quad \times \frac{2}{\theta_R}\,\mathrm{e}^{-\frac{2}{\theta_R}(t-\tau_R)}, & \text{if } t > \tau_R, \end{cases} \tag{S13}
$$

which depends on all seven parameters in $\Theta_{\text{SC}}$.

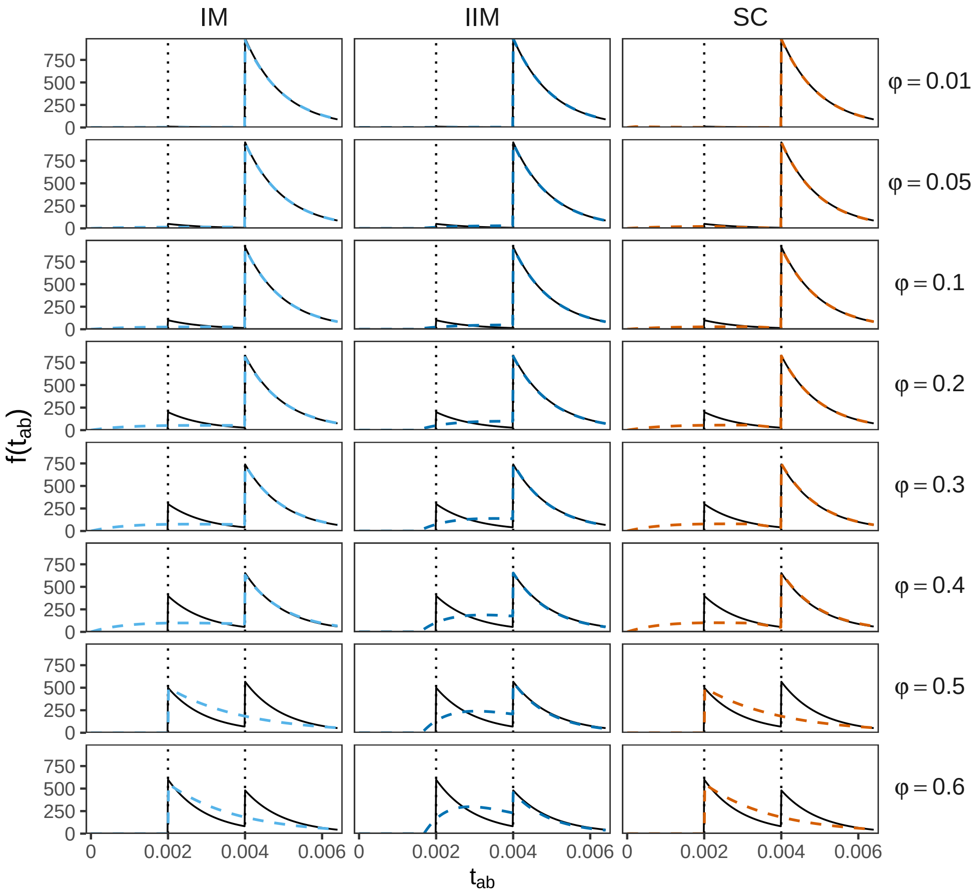


Figure S1: Distributions of the coalescent time $t_{ab}$ between a sequence from $A$ and a sequence from $B$ under models of figure 1. The true distribution under the MSC-I model of figure 1**a** for different values of introgression probability $\varphi$ is shown in solid curve, while the best-fitting distributions under the MSC-M models (IM, IIM, SC) of figure 1**b**–**d**), calculated by minimizing the KL divergence from the true MSC-I model (eq. 1) under the constraint $\theta_A = \theta_R$, are shown as dashed curves (see fig. 2).

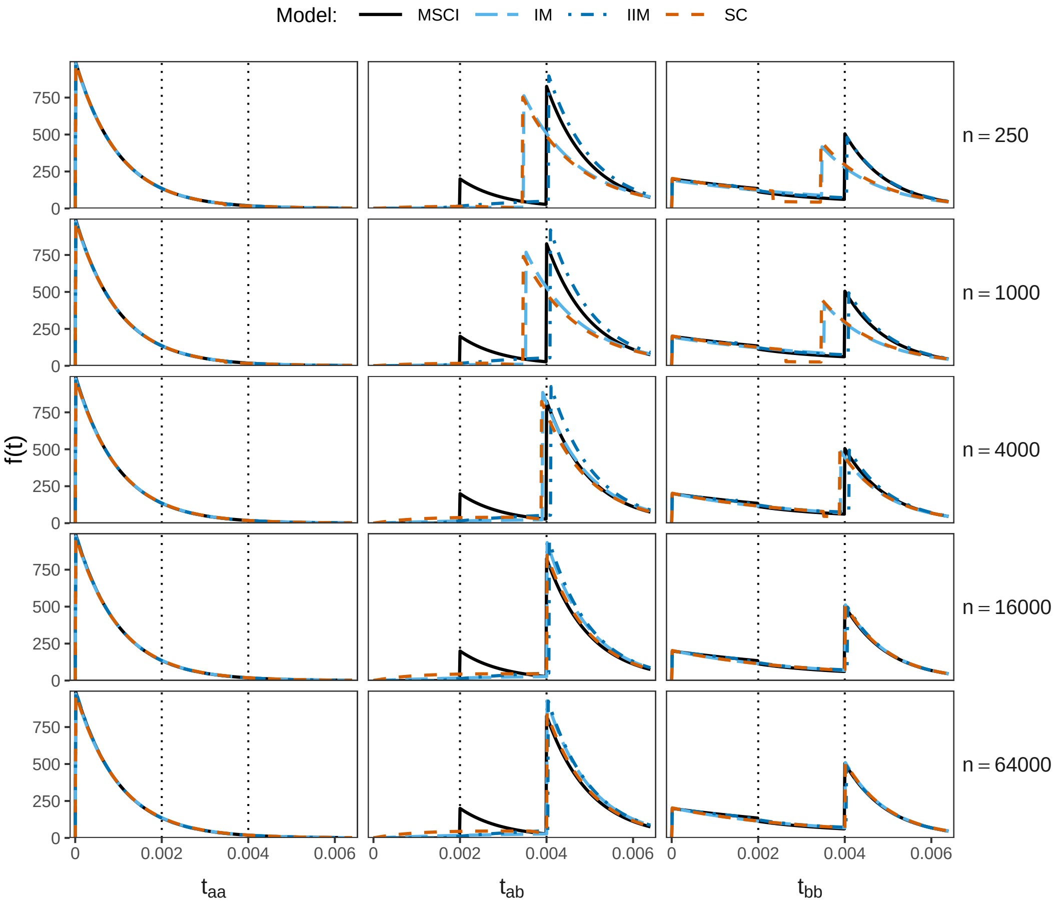


Figure S2: Distributions of coalescent times between two sequences ($t_{aa}$, $t_{ab}$ and $t_{bb}$) under the true MSC-I model (fig. 1**a**) with different numbers of sites per locus ($n$) and under the MSC-M models (IM, IIM, SC; fig. 1**b**-**d**) calculated using the best-fitting parameter values obtained from BPP analysis of data of $L = 4{,}000$ loci (fig. 3, first column). See legend to figure 4.

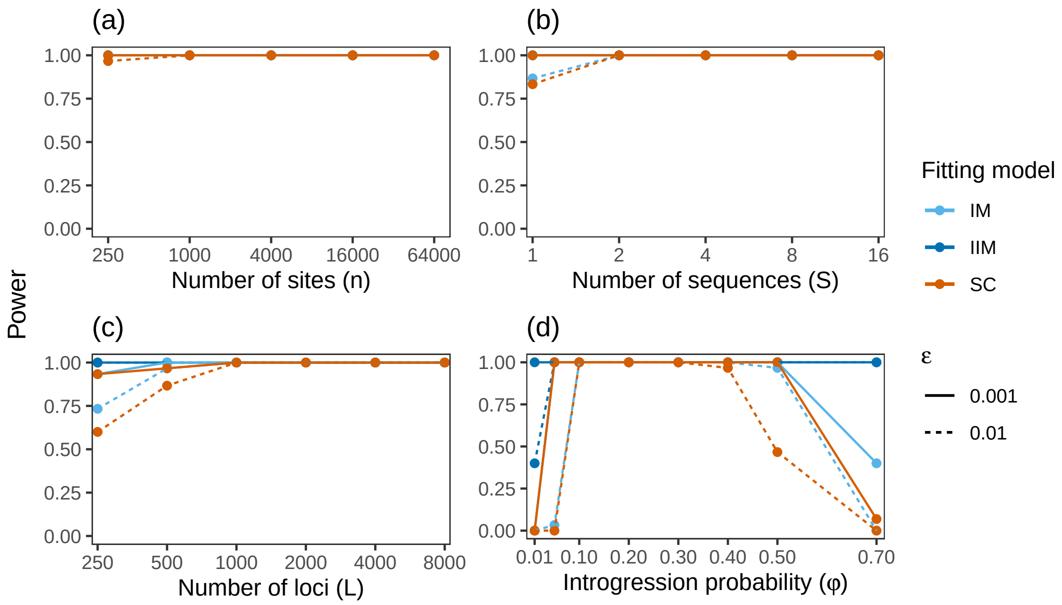


Figure S3: Power of the Bayesian test for gene flow applied to simulated data under the MSC-I model of figure 1**a**. The Bayes factor $B_{10}$ was calculated using the Savage-Dickey density ratio with two small values for null effect (no gene flow): $\varepsilon = 0.01$ and 0.001. We tested $H_0 : M_{A\to B} = 0$ against $H_1 : M_{A\to B} > 0$ under the IM, IIM or SC models (fig. 1**b**-**d**). The power was calculated as the proportion of replicate datasets in which $B_{10} > 100$. Parameter estimates are summarized in figure 3.

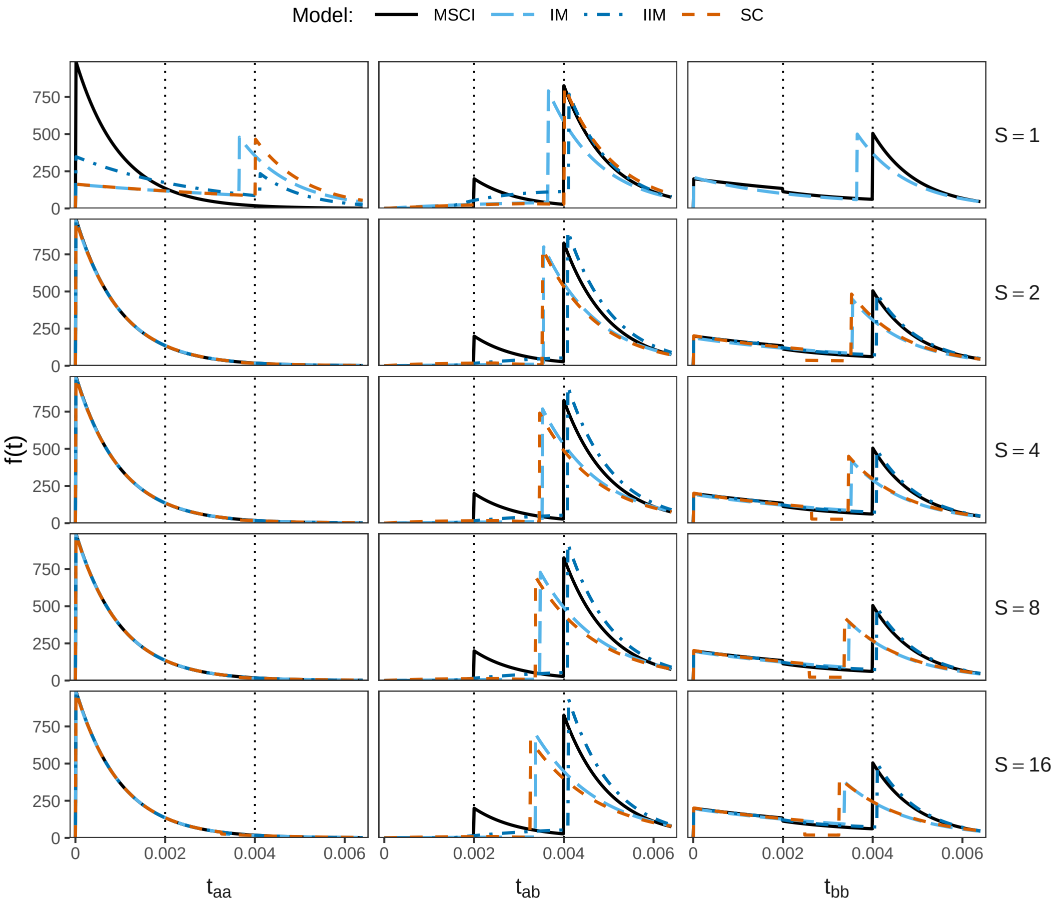


Figure S4: Distributions of coalescent times between two sequences ($t_{aa}$, $t_{ab}$ and $t_{bb}$) under the true MSC-I model (fig. 1**a**) at different numbers of sequences per species ($S$) and under the three fitting MSC-M models (IM, IIM, SC; fig. 1**b**-**d**) calculated using the best-fitting parameter values obtained from BPP analysis of data of $L = 4{,}000$ loci (fig. 3, second column).

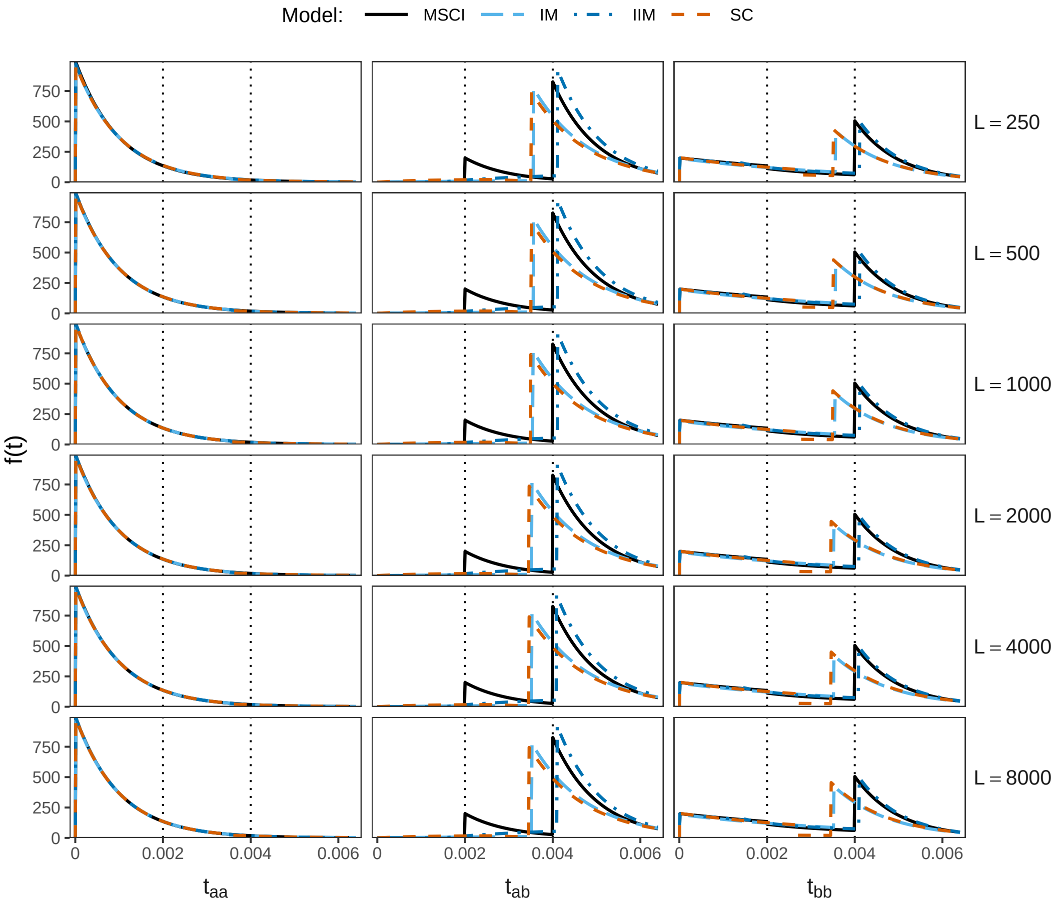


Figure S5: Distributions of coalescent times between two sequences ($t_{aa}$, $t_{ab}$ and $t_{bb}$) under the true MSC-I model (fig. 1**a**) at different numbers of loci ($L$) and under the three fitting MSC-M models (IM, IIM, SC; fig. 1**b**-**d**) using the best-fitting parameter values obtained from BPP (fig. 3, third column). Note that different choices of $L$ led to highly similar estimates and coalescent time distributions.

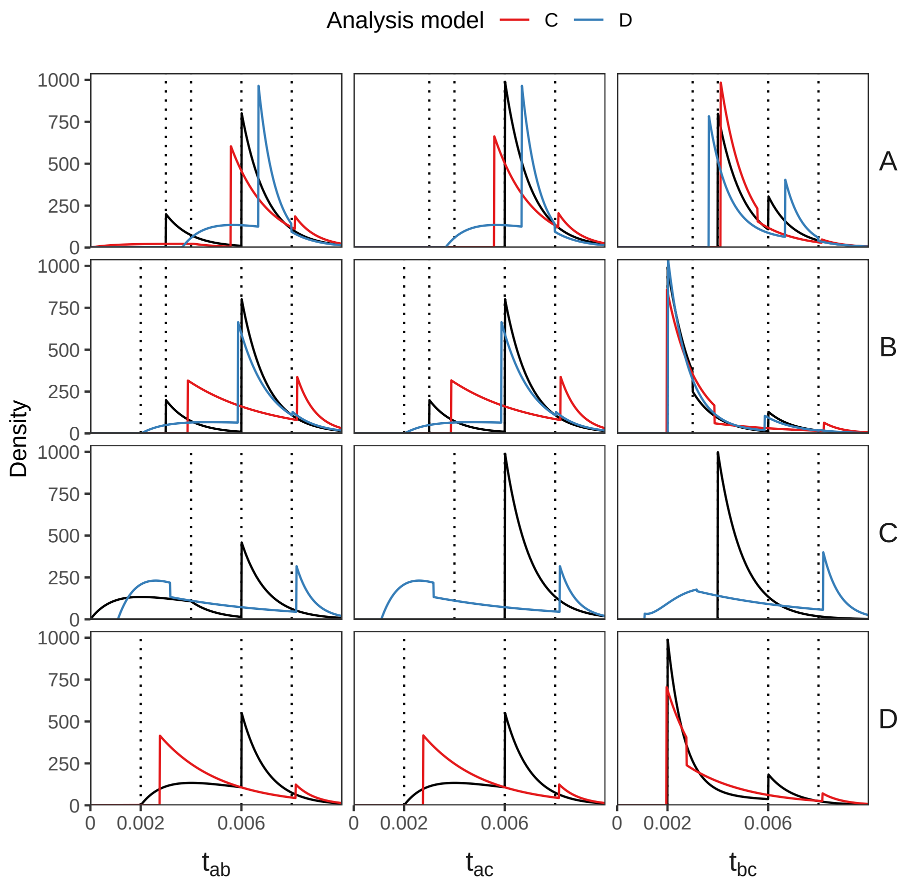


Figure S6: The true (black) and fitting (red and blue) distributions of coalescent times between sequences from two species, $f(t_{ab}), f(t_{ac}), f(t_{bc})$, when data are generated under models A, B, C, and D of figure 5 and analyzed under models C and D. Each row corresponds to a true model (A, B, C, or D). Each colour line indicates the fitting model (C or D). For example, the first row shows the A-C (red) and A-D (blue) settings (fig. 5). The fitting distributions were calculated using posterior means of parameters from BPP analysis of simulated data of $L = 4{,}000$ loci, averaged across 100 replicates (fig. 5**e**).

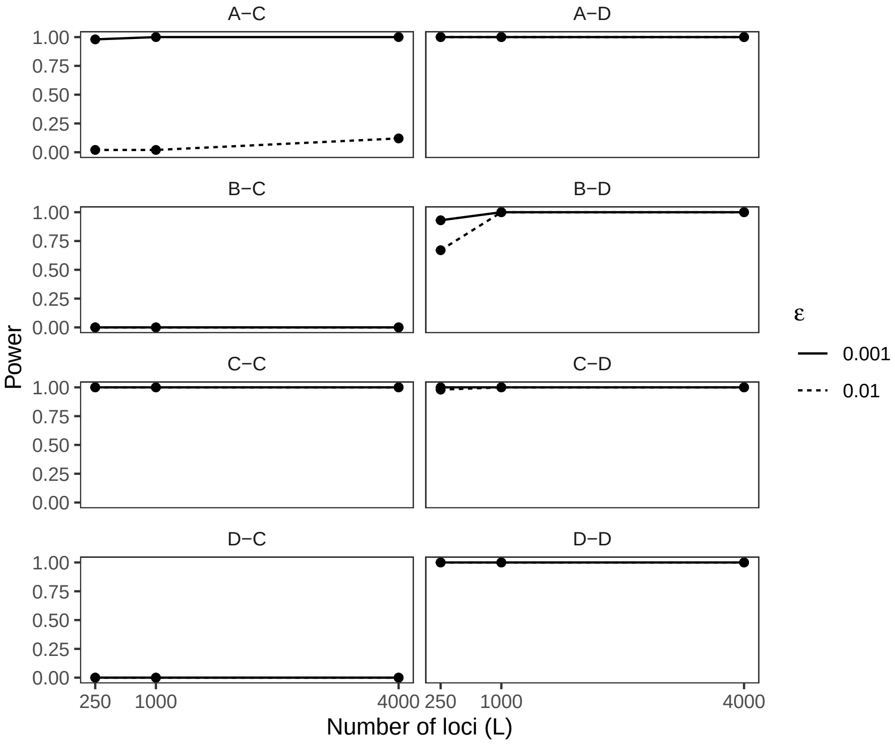


Figure S7: Power of the Bayesian test for gene flow applied to simulated data under models A-D of figure 5. We tested $H_0 : M_{A \to B} = 0$ against $H_1 : M_{A \to B} > 0$ under models C (first column) or D (second column). Parameter estimates are shown in figure 5**e**. See legend to figure S3 for more details.

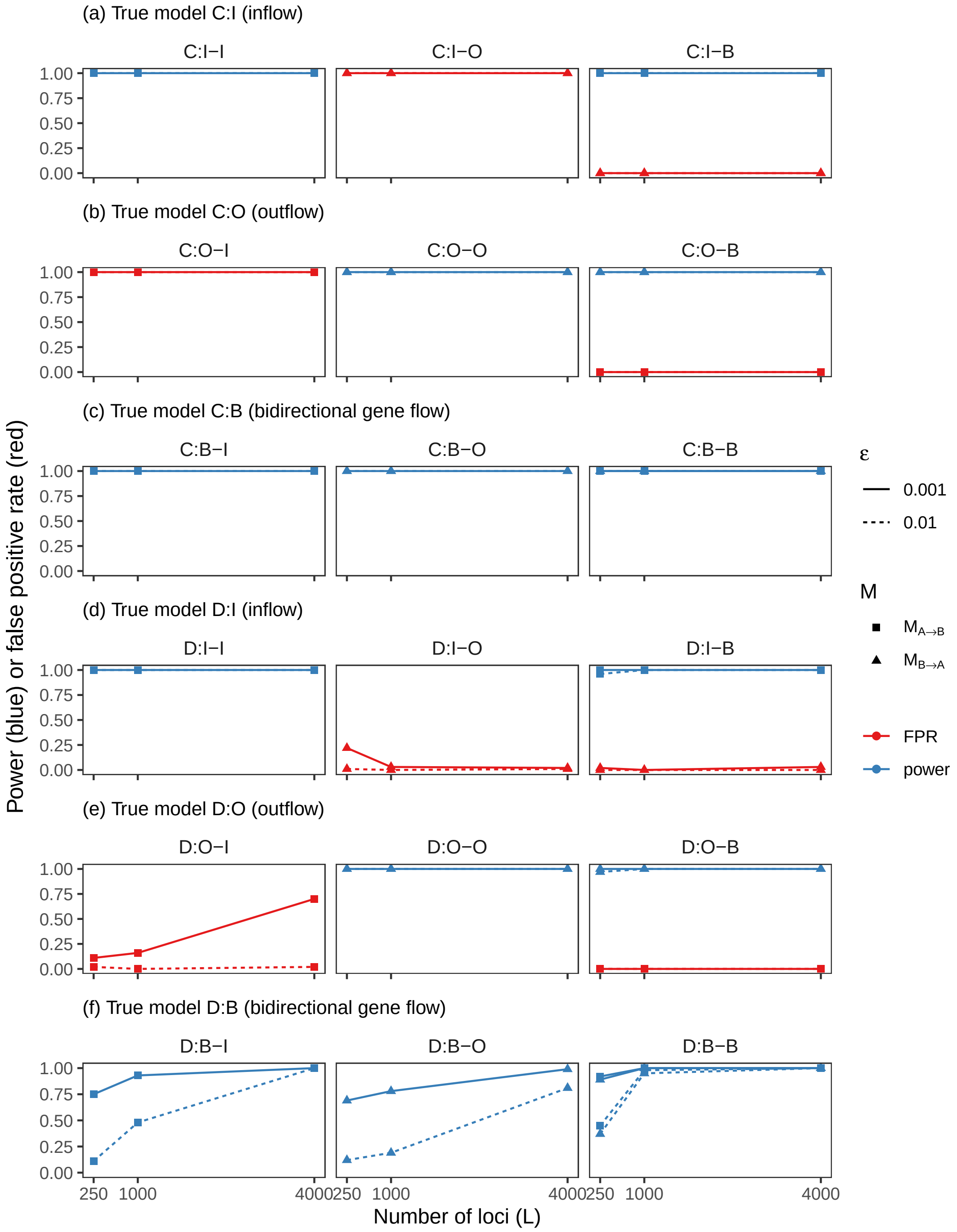


Figure S8: Power (blue) and false positive rate (FPR; red) of the Bayesian test for gene flow applied to data of four species simulated under the C and D models: (a) C:I (inflow), (b) C:O (outflow), (c) C:B (bidirectional), (d) D:I (inflow), (e) D:O (outflow), (f) D:B (bidirectional) (fig. 6**a**-**f**). The data were analyzed under I, O, and B models. Parameter estimates are shown in figure 6**g**-**h**. See legend to figure S3 for more details.

**Table S1. Posterior means and 95% HPD intervals (in parentheses) for parameters, averaged over 100 replicated datasets, obtained from BPP analysis of simulated datasets of $L = 4{,}000$ loci for the eight simulation-analysis model settings in figure 5e**

| | A-C | B-C | C-C | D-C | A-D | B-D | C-D | D-D |
|---|---|---|---|---|---|---|---|---|
| $\theta_A$ | 1.99 (1.93, 2.04) | 1.96 (1.90, 2.02) | 2.00 (1.94, 2.05) | 1.91 (1.86, 1.97) | 2.00 (1.94, 2.06) | 2.00 (1.94, 2.06) | 2.18 (2.12, 2.24) | 2.00 (1.94, 2.05) |
| $\theta_B$ | 1.92 (1.87, 1.98) | 1.99 (1.93, 2.05) | 2.01 (1.94, 2.08) | 1.98 (1.92, 2.05) | 2.06 (2.00, 2.12) | 2.01 (1.94, 2.07) | 2.61 (2.52, 2.69) | 2.00 (1.94, 2.06) |
| $\theta_C$ | 2.00 (1.94, 2.06) | 2.00 (1.94, 2.06) | 2.00 (1.94, 2.06) | 2.00 (1.94, 2.06) | 1.93 (1.88, 1.99) | 2.01 (1.94, 2.07) | 1.12 (1.09, 1.16) | 2.00 (1.94, 2.06) |
| $\theta_D$ | 2.00 (1.94, 2.06) | 1.99 (1.93, 2.05) | 2.00 (1.94, 2.06) | 1.99 (1.93, 2.05) | 2.00 (1.94, 2.06) | 2.00 (1.94, 2.06) | 1.97 (1.91, 2.03) | 2.00 (1.94, 2.05) |
| $\theta_R$ | 1.75 (1.48, 2.02) | 1.48 (1.17, 1.79) | 1.99 (1.71, 2.26) | 1.65 (1.32, 1.97) | 2.04 (1.76, 2.32) | 1.96 (1.65, 2.26) | 1.30 (0.99, 1.61) | 1.99 (1.66, 2.32) |
| $\theta_S$ | 2.99 (2.64, 3.34) | 6.32 (5.91, 6.72) | 2.03 (1.68, 2.38) | 4.77 (4.57, 4.98) | 1.36 (0.94, 1.79) | 2.36 (1.76, 2.98) | 9.29 (8.80, 9.78) | 2.03 (1.25, 2.77) |
| $\theta_T$ | 2.02 (1.72, 2.32) | 2.30 (2.15, 2.45) | 2.05 (1.58, 2.52) | 2.83 (2.50, 3.16) | 2.55 (2.24, 2.86) | 1.92 (1.78, 2.05) | 58.46 (49.94, 67.25) | 2.00 (1.85, 2.16) |
| $\tau_R$ | 8.12 (7.97, 8.27) | 8.21 (8.04, 8.38) | 8.00 (7.85, 8.15) | 8.15 (7.97, 8.33) | 7.98 (7.83, 8.13) | 8.02 (7.86, 8.19) | 8.18 (8.01, 8.35) | 8.01 (7.83, 8.18) |
| $\tau_S$ | 5.57 (5.44, 5.70) | 3.87 (3.78, 3.97) | 6.00 (5.85, 6.15) | 2.75 (2.69, 2.81) | 6.67 (6.42, 6.92) | 5.86 (5.56, 6.17) | 3.17 (3.04, 3.29) | 5.98 (5.31, 6.66) |
| $\tau_T$ | 4.11 (4.00, 4.22) | 1.97 (1.91, 2.02) | 3.99 (3.85, 4.13) | 1.96 (1.91, 2.02) | 3.65 (3.55, 3.74) | 2.02 (1.97, 2.07) | 1.09 (1.06, 1.12) | 2.00 (1.95, 2.05) |
| $M$ | 0.01 (0.01, 0.01) | 0.00 (0.00, 0.00) | 0.20 (0.19, 0.21) | 0.00 (0.00, 0.00) | 0.13 (0.12, 0.14) | 0.04 (0.04, 0.04) | 6.80 (5.75, 7.88) | 0.20 (0.19, 0.21) |

Note.— Values of $\tau$ and $\theta$ are multiplied by $10^3$. Results for all data sizes ($L = 250, 1000, 4000$ loci) are shown in figure 5**e**.

**Table S2. Posterior means and 95% HPD intervals (in parentheses) for parameters, averaged over 100 replicated datasets, obtained from BPP analysis of simulated datasets of $L = 4000$ loci for the nine simulation-analysis model settings for model C of figure 6a-c**

| | Model C:I | | | | Model C:O | | | | Model C:B | | | |
|---|---|---|---|---|---|---|---|---|---|---|---|---|
| | $\Theta_I$ | $\hat{\Theta}_I$ (C:I-I) | $\hat{\Theta}_O$ (C:I-O) | $\hat{\Theta}_B$ (C:I-B) | $\Theta_O$ | $\hat{\Theta}_I$ (C:O-I) | $\hat{\Theta}_O$ (C:O-O) | $\hat{\Theta}_B$ (C:O-B) | $\Theta_B$ | $\hat{\Theta}_I$ (C:B-I) | $\hat{\Theta}_O$ (C:B-O) | $\hat{\Theta}_B$ (C:B-B) |
| $\theta_A$ | 2.0 | 2.00 (1.94, 2.05) | 0.87 (0.83, 0.91) | 1.99 (1.93, 2.04) | 2.0 | 3.84 (3.74, 3.93) | 2.00 (1.93, 2.07) | 2.00 (1.93, 2.07) | 2.0 | 3.60 (3.51, 3.68) | 1.28 (1.22, 1.34) | 2.00 (1.91, 2.08) |
| $\theta_B$ | 2.0 | 2.01 (1.94, 2.08) | 3.73 (3.64, 3.83) | 2.01 (1.94, 2.08) | 2.0 | 0.92 (0.88, 0.95) | 2.00 (1.94, 2.06) | 1.99 (1.93, 2.05) | 2.0 | 1.29 (1.24, 1.35) | 3.43 (3.34, 3.52) | 2.00 (1.92, 2.08) |
| $\theta_C$ | 2.0 | 2.00 (1.94, 2.06) | 1.95 (1.90, 2.01) | 2.00 (1.94, 2.06) | 2.0 | 2.01 (1.95, 2.07) | 2.00 (1.94, 2.06) | 2.00 (1.95, 2.06) | 2.0 | 2.00 (1.93, 2.06) | 1.91 (1.85, 1.96) | 2.00 (1.94, 2.06) |
| $\theta_D$ | 2.0 | 2.00 (1.94, 2.06) | 2.00 (1.94, 2.06) | 2.00 (1.94, 2.06) | 2.0 | 2.00 (1.94, 2.06) | 2.00 (1.94, 2.06) | 2.00 (1.94, 2.06) | 2.0 | 1.99 (1.93, 2.05) | 2.00 (1.94, 2.06) | 2.00 (1.94, 2.06) |
| $\theta_R$ | 2.0 | 1.99 (1.71, 2.26) | 1.78 (1.48, 2.07) | 2.00 (1.72, 2.28) | 2.0 | 1.74 (1.42, 2.06) | 2.00 (1.68, 2.31) | 2.00 (1.68, 2.31) | 2.0 | 1.51 (1.19, 1.82) | 1.78 (1.47, 2.08) | 1.98 (1.68, 2.28) |
| $\theta_S$ | 2.0 | 2.03 (1.68, 2.38) | 1.81 (0.95, 2.67) | 1.99 (1.63, 2.34) | 2.0 | 3.57 (3.37, 3.78) | 2.01 (1.54, 2.49) | 2.02 (1.55, 2.50) | 2.0 | 5.26 (4.95, 5.57) | 3.27 (2.73, 3.78) | 2.08 (1.56, 2.60) |
| $\theta_T$ | 2.0 | 2.05 (1.58, 2.52) | 9.40 (8.12, 10.78) | 2.05 (1.59, 2.53) | 2.0 | 0.45 (0.14, 0.76) | 2.02 (1.79, 2.25) | 1.98 (1.74, 2.21) | 2.0 | 0.33 (0.06, 0.63) | 16.91 (10.24, 25.32) | 2.04 (1.60, 2.50) |
| $\tau_R$ | 8.0 | 8.00 (7.85, 8.15) | 8.09 (7.94, 8.25) | 8.00 (7.85, 8.15) | 8.0 | 8.11 (7.94, 8.29) | 8.01 (7.84, 8.17) | 8.01 (7.84, 8.17) | 8.0 | 8.18 (8.01, 8.35) | 8.09 (7.93, 8.25) | 8.01 (7.85, 8.17) |
| $\tau_S$ | 6.0 | 6.00 (5.85, 6.15) | 6.80 (6.30, 7.30) | 6.02 (5.87, 6.18) | 6.0 | 3.73 (3.65, 3.80) | 6.00 (5.75, 6.25) | 6.00 (5.75, 6.25) | 6.0 | 3.76 (3.68, 3.85) | 4.89 (4.48, 5.34) | 5.95 (5.66, 6.26) |
| $\tau_T$ | 4.0 | 3.99 (3.85, 4.13) | 4.08 (3.97, 4.18) | 3.99 (3.85, 4.13) | 4.0 | 3.49 (3.34, 3.63) | 3.99 (3.91, 4.08) | 4.01 (3.92, 4.10) | 4.0 | 3.58 (3.42, 3.73) | 3.33 (3.22, 3.43) | 3.99 (3.86, 4.11) |
| $M_{A \to B}$ | 0.2 | 0.20 (0.19, 0.21) | n/a | 0.20 (0.19, 0.21) | n/a | 0.17 (0.16, 0.17) | n/a | 0.00 (0.00, 0.00) | 0.2 | 0.34 (0.33, 0.36) | n/a | 0.20 (0.19, 0.21) |
| $M_{B \to A}$ | n/a | n/a | 0.55 (0.51, 0.59) | 0.01 (0.00, 0.02) | 0.2 | n/a | 0.20 (0.19, 0.21) | 0.20 (0.19, 0.21) | 0.2 | n/a | 0.35 (0.33, 0.37) | 0.20 (0.19, 0.21) |

Note.— Values of $\tau$ and $\theta$ are multiplied by $10^3$. Results for all data sizes ($L = 250, 1000, 4000$ loci) are shown in figure 6**g**. 'n/a' indicates that the parameter does not exist in a given model.

**Table S3. Posterior means and 95% HPD intervals (in parentheses) for parameters, averaged over 100 replicated datasets, obtained from BPP analysis of simulated datasets of $L = 4000$ loci for the nine simulation-analysis model settings for model D of figure 6d-f**

| | Model D:I | | | | Model D:O | | | | Model D:B | | | |
|---|---|---|---|---|---|---|---|---|---|---|---|---|
| | $\Theta_I$ | $\hat{\Theta}_I$ (D:I-I) | $\hat{\Theta}_O$ (D:I-O) | $\hat{\Theta}_B$ (D:I-B) | $\Theta_O$ | $\hat{\Theta}_I$ (D:O-I) | $\hat{\Theta}_O$ (D:O-O) | $\hat{\Theta}_B$ (D:O-B) | $\Theta_B$ | $\hat{\Theta}_I$ (D:B-I) | $\hat{\Theta}_O$ (D:B-O) | $\hat{\Theta}_B$ (D:B-B) |
| $\theta_A$ | 2.0 | 2.00 (1.94, 2.05) | 1.91 (1.85, 1.97) | 1.99 (1.93, 2.04) | 2.0 | 2.12 (2.06, 2.18) | 2.00 (1.94, 2.06) | 2.00 (1.95, 2.06) | 2.0 | 2.05 (1.99, 2.11) | 1.98 (1.92, 2.04) | 2.00 (1.94, 2.06) |
| $\theta_B$ | 2.0 | 2.00 (1.94, 2.06) | 2.00 (1.94, 2.06) | 2.00 (1.94, 2.06) | 2.0 | 2.02 (1.96, 2.08) | 2.00 (1.94, 2.06) | 2.01 (1.94, 2.07) | 2.0 | 1.99 (1.93, 2.06) | 1.99 (1.93, 2.05) | 1.99 (1.93, 2.06) |
| $\theta_C$ | 2.0 | 2.00 (1.94, 2.06) | 2.00 (1.93, 2.06) | 2.00 (1.94, 2.07) | 2.0 | 2.02 (1.96, 2.08) | 2.00 (1.94, 2.06) | 2.00 (1.94, 2.06) | 2.0 | 2.00 (1.94, 2.06) | 1.99 (1.93, 2.06) | 2.00 (1.94, 2.06) |
| $\theta_D$ | 2.0 | 2.00 (1.94, 2.05) | 2.00 (1.94, 2.06) | 2.00 (1.94, 2.06) | 2.0 | 2.00 (1.94, 2.06) | 2.00 (1.95, 2.06) | 2.00 (1.95, 2.06) | 2.0 | 2.00 (1.94, 2.06) | 2.00 (1.94, 2.06) | 2.00 (1.94, 2.06) |
| $\theta_R$ | 2.0 | 1.99 (1.66, 2.32) | 1.59 (1.24, 1.92) | 2.00 (1.67, 2.33) | 2.0 | 1.80 (1.47, 2.12) | 1.99 (1.65, 2.32) | 2.00 (1.67, 2.33) | 2.0 | 1.68 (1.34, 2.02) | 1.69 (1.35, 2.02) | 1.98 (1.65, 2.31) |
| $\theta_S$ | 2.0 | 2.03 (1.25, 2.77) | 4.80 (4.59, 5.00) | 2.00 (1.25, 2.73) | 2.0 | 4.01 (3.80, 4.22) | 1.92 (1.17, 2.63) | 1.81 (1.06, 2.52) | 2.0 | 4.59 (4.37, 4.81) | 4.44 (4.13, 4.72) | 2.16 (1.34, 2.96) |
| $\theta_T$ | 2.0 | 2.00 (1.85, 2.16) | 2.86 (2.52, 3.19) | 2.01 (1.85, 2.17) | 2.0 | 1.25 (1.11, 1.39) | 2.00 (1.90, 2.10) | 1.94 (1.82, 2.05) | 2.0 | 1.86 (1.59, 2.14) | 2.69 (2.26, 3.12) | 2.04 (1.85, 2.25) |
| $\tau_R$ | 8.0 | 8.01 (7.83, 8.18) | 8.18 (8.00, 8.36) | 8.00 (7.83, 8.17) | 8.0 | 8.08 (7.90, 8.25) | 8.00 (7.83, 8.18) | 8.00 (7.82, 8.17) | 8.0 | 8.14 (7.96, 8.32) | 8.14 (7.96, 8.32) | 8.01 (7.84, 8.19) |
| $\tau_S$ | 6.0 | 5.98 (5.31, 6.66) | 2.76 (2.69, 2.83) | 5.98 (5.34, 6.63) | 6.0 | 3.08 (2.99, 3.17) | 6.07 (5.44, 6.72) | 6.13 (5.51, 6.78) | 6.0 | 2.82 (2.67, 2.98) | 3.04 (2.78, 3.37) | 5.85 (5.14, 6.56) |
| $\tau_T$ | 2.0 | 2.00 (1.95, 2.05) | 1.96 (1.90, 2.02) | 2.00 (1.95, 2.05) | 2.0 | 2.10 (2.05, 2.16) | 2.00 (1.96, 2.05) | 2.01 (1.97, 2.06) | 2.0 | 1.99 (1.94, 2.05) | 1.96 (1.90, 2.02) | 1.99 (1.95, 2.04) |
| $M_{A \to B}$ | 0.2 | 0.20 (0.19, 0.21) | n/a | 0.20 (0.19, 0.21) | n/a | 0.02 (0.01, 0.03) | n/a | 0.00 (0.00, 0.01) | 0.2 | 0.19 (0.15, 0.23) | n/a | 0.20 (0.17, 0.22) |
| $M_{B \to A}$ | n/a | n/a | 0.02 (0.00, 0.04) | 0.01 (0.00, 0.01) | 0.2 | n/a | 0.20 (0.19, 0.22) | 0.20 (0.18, 0.21) | 0.2 | n/a | 0.25 (0.14, 0.35) | 0.20 (0.16, 0.24) |

Note.— Values of $\tau$ and $\theta$ are multiplied by $10^3$. Results for all data sizes ($L = 250, 1000, 4000$ loci) are shown in figure 6**h**. 'n/a' indicates that the parameter does not exist in a given model.
